# A combinatorial indexing strategy for epigenomic profiling of plant single cells

**DOI:** 10.1101/2021.11.07.467612

**Authors:** Xiaoyu Tu, Alexandre P. Marand, Robert J. Schmitz, Silin Zhong

**Affiliations:** The South China Botanical Garden, Chinese Academy of Sciences, Guangzhou, China; State Key Laboratory of Agrobiotechnology, School of Life Sciences, The Chinese University of Hong Kong, Hong Kong, China; Department of Genetics, University of Georgia, Athens, GA 30602, USA

**Keywords:** combinatorial indexing, single-cell ATAC-seq, epigenome

## Abstract

Understanding how *cis-*regulatory elements facilitate gene expression is a key question in biology. Recent advances in single-cell genomics have led to the discovery of cell-specific chromatin landscapes that underlie transcription programs. However, the high equipment and reagent costs of commercial systems limit their applications for many laboratories. In this study, we profiled the Arabidopsis root single-cell epigenome using a combinatorial index and dual PCR barcode strategy without the need of any specialized equipment. We generated chromatin accessibility profiles for 13,576 *Arabidopsis thaliana* root nuclei with an average of 12,784 unique Tn5 integrations per cell and 85% of the Tn5 insertions localizing to discrete accessible chromatin regions. Comparison with data generated from a commercial microfluidic platform revealed that our method is capable of unbiased identification of cell type-specific chromatin accessibility with improved throughput, quality, and efficiency. We anticipate that by removing cost, instrument, and other technical obstacles, this combinatorial indexing method will be a valuable tool for routine investigation of single-cell epigenomes and usher new insight into plant growth, development and their interactions with the environment.

## INTRODUCTION

In multicellular organisms, each cell shares an identical set of genetic instructions despite having highly specialized structures and functions. Cell-specific physiology due to dynamic gene expression characteristics are often the result of the binding and activity of cell-specific transcription factors (TFs) to proximal and distal *cis*-acting regulatory elements (CREs) in accessible chromatin regions (ACRs) associated with a given gene (Kundaje et al., 2015; Thurman et al., 2012). Chromatin accessibility profiling methods such as DNase I hypersensitive site sequencing (DNase-seq) and Assay for Transposase Accessible Chromatin sequencing (ATAC-seq) have been developed to measure the chromatin accessibility as a generalized proxy for the regulatory DNA across numerous plant species (Buenrostro et al., 2013; Crawford et al., 2006; Minnoye et al., 2021; Moore et al., 2020). However, these assays can only capture the average chromatin accessibility signal across a population of cells, masking cell-specific and rare events within a given tissue.

Recent development of droplet microfluidic device-based single-cell systems have enabled researchers to co-encapsulate individual cells or nuclei with barcode beads in nanoliter water-in-oil emulsions, making it possible to perform sequencing at the single-cell level (Lareau et al., 2019; Satpathy et al., 2019). For example, single-cell ATAC-seq (scATAC-seq) assays based on the 10X Genomics Chromium system have been successfully applied to study cell-specific chromatin accessibility in plants (Dorrity et al., 2021; Farmer et al., 2021; Marand et al., 2021; Marand and Schmitz, 2022). However, these commercial single-cell assays often require specialized equipment, trained personnel, and are very expensive, making it difficult for an individual laboratory to perform in house. To bypass the need of commercial microfluidic systems, single-cell ATAC-seq based on the combinatorial indexing strategy (sci-ATAC-seq) has been developed and widely used for animal research (Cusanovich et al., 2015). Such easy to use, low-cost, sensitive and high-throughput epigenomic profiling methods are needed for plant systems to decode the full repertoire of *cis*-regulatory diversity and chromatin dynamics in different tissues throughout the life cycle and in response to environmental stimuli.

To meet this challenge, we developed an inexpensive and simple single-cell ATAC-seq method based on combinatorial indexing and dual PCR barcoding, without the need of any specialized equipment. We applied it to *Arabidopsis thaliana* root tissues and benchmarked our single-cell chromatin accessibility profiles to previously generated 10X Genomics scATAC-seq data from the same tissue. From two biological replicates, we obtained high-quality chromatin accessibility data for 13,576 cells. Comparison with published data sets generated by the commercial 10X systems revealed that sci-ATAC-seq readily captures known cell-type-specific chromatin accessibility profiles, is similarly correlated with nuclear transcription, and is generalized by lower background signal, organellar contamination, and doublet rates. We anticipate that this technological breakthrough will empower plant researchers to overcome the difficulty and cost barriers of single-cell epigenome studies, enabling routine single-cell epigenomic profiling at a larger scale.

## RESULTS

### Implementation of sci-ATAC-seq to profile single-nucleus chromatin accessibility

To demonstrate the utility of combinatorial indexing to resolve heterogeneous chromatin accessibility signals from complex plant tissues, we applied it to whole root tissues from 2-week-old *A. thaliana* plants (**Figure 1 and Supplementary information**). First, isolated nuclei were split into four 96-well plates with 24×16 barcoded Tn5 transposon to integrate 384 adapter combinations into the accessible chromatin regions (ACRs) of individual nuclei. The tagmentation reaction is then quenched and tagged nuclei are pooled and redistributed to multiple 96-wells plates with approximately 15 nuclei per well. Plate PCR was then performed using 12×8 sets of unique dual-indexed primers complementary to the transposase-introduced adapters, which adds a second round of well-specific barcodes to Tn5-tagged DNA. Subsequently, PCR products from individual wells in each plate are combined and purified, and a final round of PCR is performed to add a third barcode into each molecule to label the sample plate. In this method, a single 96-well plate can generate sci-ATAC-seq profiles for ∼1,500 cells, and one could easily perform 10-20 plate PCRs to obtain 15,000-30,000 chromatin accessibility profiles, making this method highly scalable.

**Figure 1.**
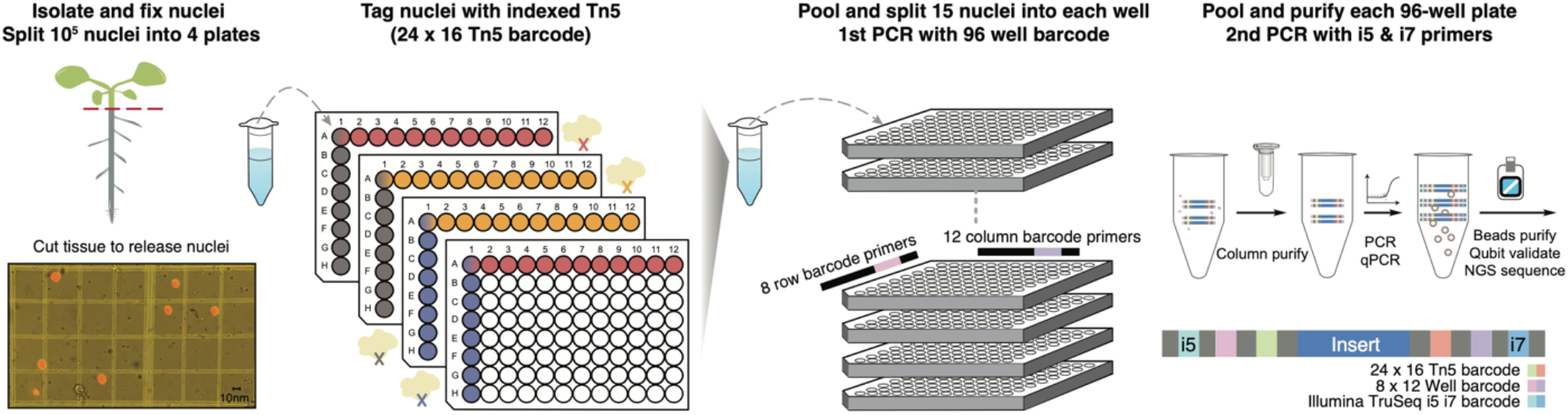
Overview of single-cell combinatorial indexing ATAC-seq. Schematic illustration of the sci-ATAC-seq experimental workflow. Tagmentation of permeabilized nuclei was carried out using the 384 barcoded Tn5 in four 96-well plates. After the first pooling, ∼15 nuclei were split into each well, and PCR was performed to introduce the second set of barcodes. Finally, PCR products from the same plate were pooled and purified for the final amplification with the third set of barcodes.

### Comparison of sci-ATAC-seq and scATAC-seq

To evaluate the efficacy of our protocol, we compared our sci-ATAC-seq data with two recent *A. thaliana* root scATAC-seq data sets generated using the commercial 10X Genomics Chromium platform (Farmer et al., 2021; Marand et al., 2021). We uniformly processed all libraries to de-multiplex reads, assign cell barcodes, align fragments to the TAIR10 reference genome, remove duplicated fragments, and use the R package *Socrates* to analyze all three data sets (**Methods**).

We first evaluated several common quality metrics, including organelle DNA contamination, doublet rate, fragment size, unique Tn5 insertion per cell and proportion of insertions within promoter and ACRs (**Figure 2**). The sci-ATAC-seq data showed lower organelle DNA contaminations when compared to the published ones generated using the commercial 10X Genomics system (**Figure 2A-B**). All libraries were enriched in subnucleosomal fragments, while the double rate and number of insertions per cell were comparable between the sci-ATAC-seq and the 10X data (**Figure 2C-E**). We also observed increased Tn5 insertions mapping to gene promoter region and within ACRs in the sci-ATAC-seq data (**Figure 2F-G**). Importantly, these gains in data quality did not come at the cost of reduced throughput. The total number of nuclei passing quality control in the sci-ATAC-seq libraries were similar to previously published 10X scATAC-seq data with a greater proportion of recovery (∼90% vs 65%) (**Figure 2H; Supplementary Figure 1**) Overall, our analysis showed that a sci-ATAC-seq experiment with only ten 96-well-plate are sufficient to generate high-quality epigenome profile for 13,576 nuclei with an average of 12,784 unique Tn5 integrations per cell and 85% of Tn5 insertions localizing to discrete ACRs. Analysis of both aggregated and single-cell signals indicated that sci-ATAC-seq can produce chromatin accessibility profiles equivalent to those generated by the 10X Genomics Chromium system with similar quality and reproducibility (**Figure 3A**).

**Figure 2.**
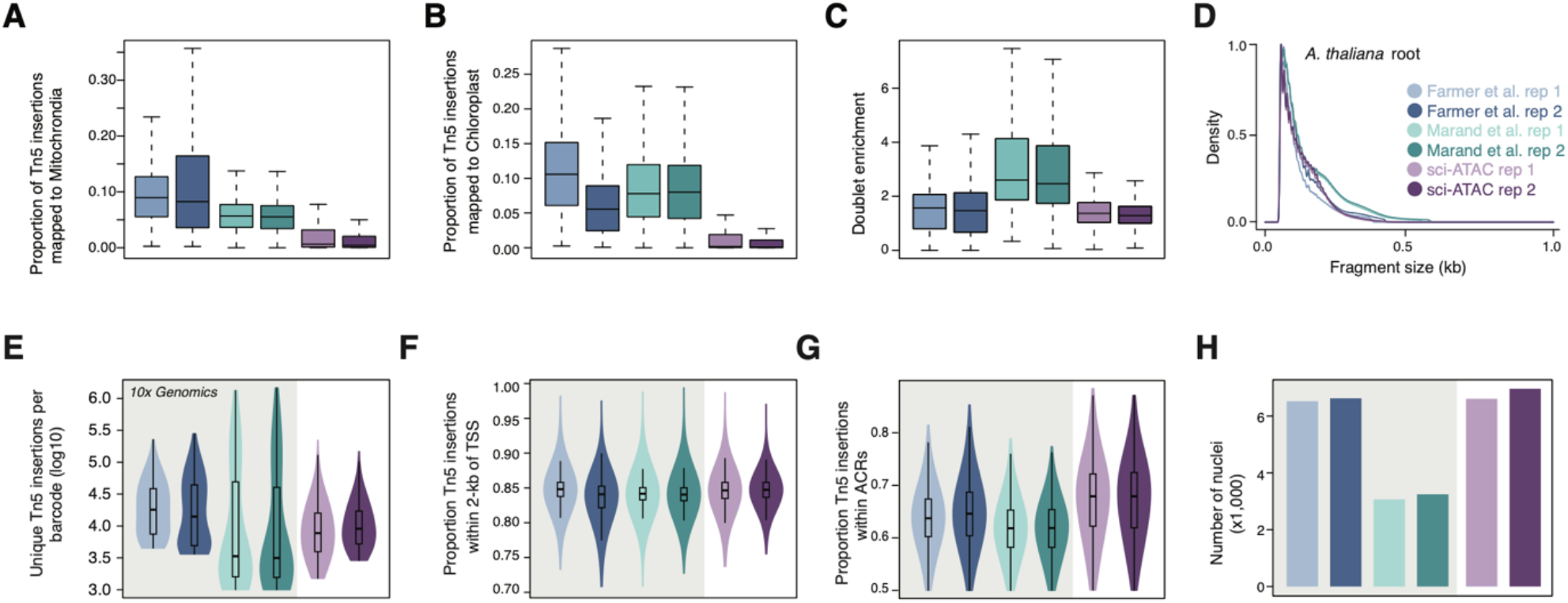
Comparison of sci-ATAC-seq quality control metrics to scATAC-seq. **(A)** Proportion of Tn5 insertions derived from the mitochondrial genome. **(B)** Proportion of Tn5 insertions derived from the chloroplast genome. **(C)** Distributions of doublet enrichment scores. **(D)** Fragment size distributions by data set and replicate. **(E)** Distribution of unique Tn5 integration sites per nucleus. **(F)** Distributions of proportion Tn5 integration sites within the promoter region 2-kb upstream of gene TSSs. **(G)** Distributions of proportion Tn5 integration sites within ACRs per nucleus. **(H)** Number of nuclei passing quality control thresholds.

**Figure 3.**
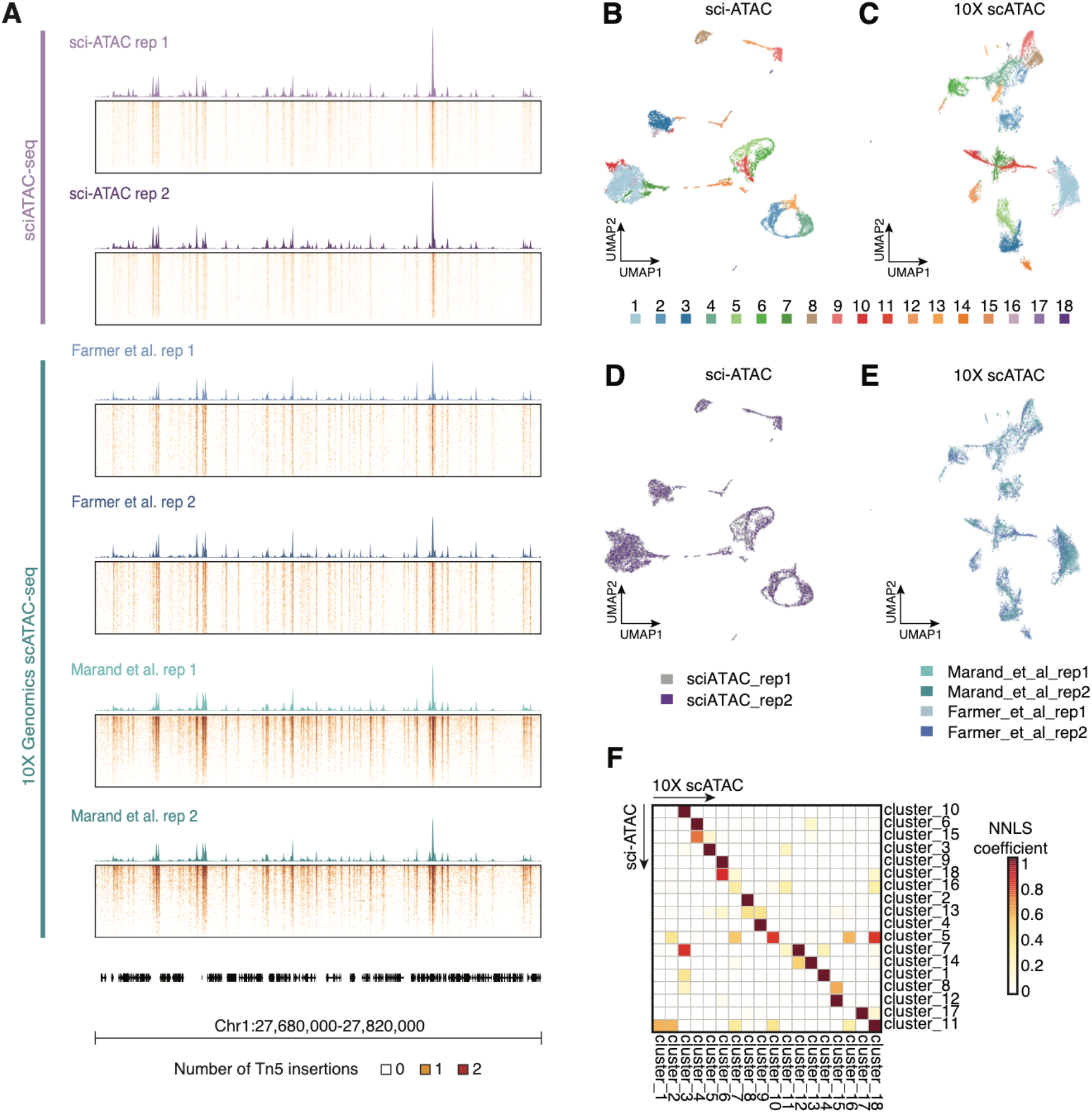
Comparison of cluster separation and replicate overlap of sci-ATAC-seq and scATAC-seq data. **(A)** Pseudobulk aggregates and single-cell accessibility profiles from 1,000 random nuclei across all data sets used in this study. **(B)** sci-ATAC-seq and **(C)** 10x scATAC-seq UMAP colored by Louvain clusters. **(D)** sci-ATAC-seq and **(E)** 10x scATAC-seq UMAP colored by biological replicate. (**F**) Non-negative least squares (NNLS) regression coefficients (described by Domcke et al. 2020. *Science*) between sci-ATAC-seq and 10X scATAC-seq data sets.

Next, we processed our sci-ATAC-seq and 10X scATAC-seq data sets in parallel to evaluate if the differences in data quality could extend to nuclei clustering. Following normalization, graph-based clustering, and UMAP visualization with *Socrates*, we identified 18 clusters in both sci-ATAC-seq and 10X scATAC-seq data sets (**Figure 3B, 3C**). Qualitative assessment of the UMAP embedding indicated that the sci-ATAC-seq replicates were highly reproducible (**Figure 3D**). A similar degree of reproducibility was observed for the 10X scATAC-seq data, albeit after first removing lab-specific batch effects (**Figure 3E**).

To identify clusters with matching chromatin accessibility profiles between data sets, we performed reciprocal non-negative least squares (NNLS) regression for each sci-ATAC-seq cluster using all 10x scATAC-seq clusters as the explanatory variable, and vice-versa, multiplying the resulting coefficients (**Methods**). NNLS analysis indicated that the cluster-aggregate chromatin accessibility profiles were well matched between sci-ATAC-seq and 10x scATAC-seq, with more than 66% of clusters uniquely matching (12/18) (**Figure 3F**). Taken together, our method provides an efficient and low-cost alternative for high-quality and high-resolution profiling of plant chromatin accessibility across tens of thousands of single nuclei with similar or better data quality compared to available commercial solutions.

### Integrative analysis of single-cell ATAC-seq and RNA-seq revealed cellular heterogeneity in the *A. thaliana* root

Next, we sought to resolve the chromatin landscapes of individual cell types present within the *A. thaliana* root. To do so, we generated a chromatin accessibility and nuclear transcription unified co-embedding by integrating three single-cell ATAC-seq and one single-nucleus RNA-seq (snRNA-seq) data set from *A. thaliana* roots using a combination of integrative non-negative matrix factorization (iNMF) and uniform manifold approximation and projection (UMAP). The co-embedding discriminated ∼1 billion pseudobulk Tn5 integration sites into 24 clusters based on shared cell states, ranging in size from 208 to 8,755 nuclei (**Figure 4A and B**). Automated annotation of cell identities using established marker genes to the patterns of nuclear transcription and chromatin accessibility recovered nearly all expected cell types of the *A. thaliana* root (**Supplementary Table 1**).

**Figure 4.**
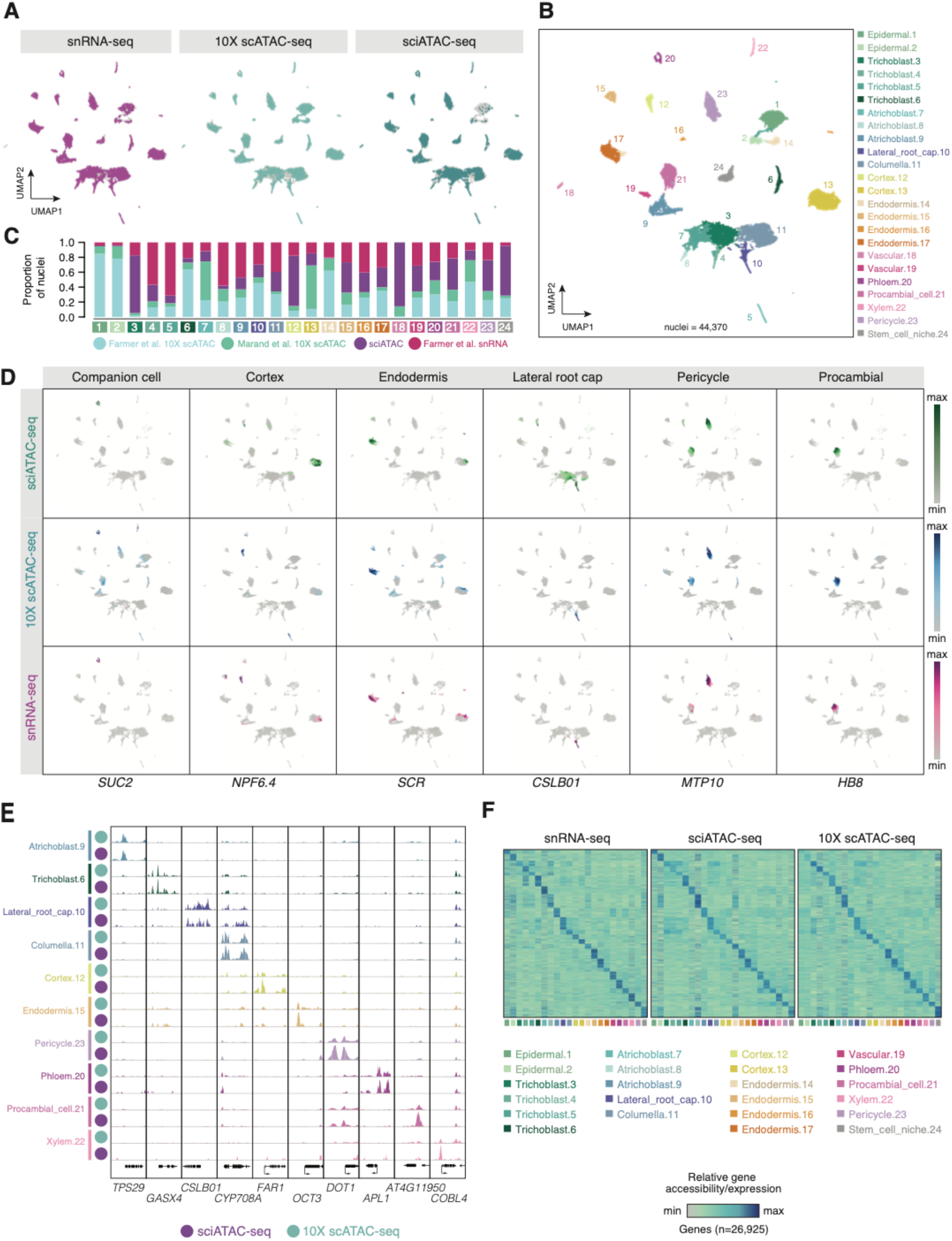
Chromatin accessibility and nuclear transcription hallmarks of *A. thaliana* root cells. **(A)** Comparison of snRNA-seq, 10x Genomics scATAC-seq, and sci-ATAC-seq from *A. thaliana* root nuclei coordinates on an integrated UMAP embedding. Grey dots indicate nuclei from a mutually exclusive dataset. **(B)** Cell-type identities of nuclei from the integrated UMAP embedding. **(C)** Proportion of nuclei derived from published 10x Genomics scATAC-seq and snRNA-seq data, as well as our sci-ATAC-seq, and sets for each cell-type cluster. **(D)** Gene accessibility and RNA abundance distributions for six cell type-specific marker genes among sci-ATAC-seq, 10x Genomics scATAC-seq, and snRNA-seq data sets. **(E)** 10x Genomics scATAC-seq and sci-ATAC-seq pseudobulk cell type Tn5 integration site coverages at 10 cell type-specific marker gene loci. Track heights denote read coverage scaled per million. **(F)** Comparison of relative snRNA-seq gene expression (left), and sci-ATAC-seq (middle) and 10x Genomics scATAC-seq (right) gene accessibility across *A. thaliana* root cell types and ∼27,000 protein coding genes.

The improved resolution and coverage of the combined data allowed us to distinguish putative cell identities by evaluating differential chromatin accessibility profiles of known cell-type-specific marker genes. For example, a known companion cell-specific *SUCROSE-PROTON SYMPORTER 2* (*SUC2*, AT1G22710) gene was found to be highly expressed in phloem cells (cluster 20) (Rodriguez-Villalon et al., 2014), concordant with elevated promoter and gene body chromatin accessibility detected in our data. Gene body chromatin accessibility for *NITRATE PEPTIDE TRANSPORTER 6*.*4* (*NPF6*.*4*, AT3G21670), a cortical parenchyma marker gene(Farmer et al., 2021), was enriched in predicted cortex clusters (cluster 12 and 13). The *SCARECROW* (*SCR*, AT3G54220) gene, which encodes a putative transcription factor that is first expressed in quiescent center precursor cells during embryogenesis (Di Laurenzio et al., 1996) and extends to the initial cells for the ground tissue (cortex and endodermis) and the endodermis (Wysocka-Diller et al., 2000), has enriched chromatin accessibility in both cortex (cluster 13) and endodermis (cluster 17) annotated clusters in our sci-ATAC-seq profiles.

To better annotate the 24 identified cell populations, we also generated pseudobulk chromatin accessibility maps for each cell-type cluster and examined the ACRs neighboring known cell-type-specific marker genes. For instance, enrichment of chromatin accessibility in *TERPENE SYNTHASE 29* (*TPS29*, AT1G31950) from cluster 9 and *XYLAN GLYCURONOSYLTRANSFERASE 4* (*GUX4*, AT1G54940) from cluster 6 suggested that the two clusters correspond to two cell types of the root epidermis, trichoblast and atrichoblast, respectively. Moreover, we observed cluster 10 with specific chromatin accessibility at the *CELLULOSE SYNTHASE-LIKE PROTEIN B1* (*CslB1*, AT2G32610) gene, a known early non hair/lateral root cap marker (Jean-Baptiste et al., 2019); cluster 11 demonstrated chromatin accessibility at *cis*-elements neighboring the *CYTOCHROME P450 CYP708 A2* (AT5G48000) gene associated with columella-specific expression (Brady et al., 2007); cluster 12 with significant enrichment at *FATTY ACID REDUCTASE 1* (*FAR1*, AT5G22500), a predicted mobile RNA associated with suberization of endodermal cells (Domergue et al., 2010); cluster 15 with enhanced chromatin accessibility at *ORGANIC CATION TRANSPORTER 3* (*OCT3*, AT1G16390), a known endodermis marker gene (Jean-Baptiste et al., 2019); ACRs significantly enriched in cluster 23 and to a lesser extent in cluster 21 at the gene locus of *DEFECTIVELY ORGANIZED TRIBUTARIES 1* (*DOT1*, AT2G36120) involved in vascular patterning; cluster 20 with enrichment of chromatin accessibility in *ALTERED PHLOEM DEVELOPMENT* (*APL*, AT1G79430), the marker gene known to appear earliest during companion cell and phloem sieve element specification (Jean-Baptiste et al., 2019; Shulse et al., 2019); cluster 21 with ACRs neighboring a putative transmembrane protein preferentially expressed in procambial cells (AT4G11950) (Brady et al., 2007); and finally differentially accessible chromatin upstream of *COBRA-LIKE 4* (*COBL4*, AT5G15630) in cluster 22, a reported marker of xylem (**Figure 4E**) (Denyer et al., 2019).

To evaluate the integrated embedding-based cell-type annotations on the independent clustering analyses of sci-ATAC-seq and 10X scATAC-seq, we plotted the inferred cell identities from the integrated analysis on to the UMAP embeddings for each scATAC-seq data set clustered in isolation. Surprisingly, we observed greater Adjusted Rand Index scores (a measure of similarity between clustering) for sci-ATAC-seq clusters relative to 10X scATAC-seq, indicating that sci-ATAC-seq data faithfully groups by cell identity independent of external data (**Supplementary Figure 3**). In addition to annotating cell types based on the aggregate chromatin profiles, we also sought to evaluate cell-type variation among modalities. We subset nuclei by snRNA-seq, 10X scATAC-seq, and sci-ATAC-seq and generated heat maps of gene expression or chromatin accessibility across 26,925 protein coding genes for each cell type (**Figure 4F**). Matched cell type comparisons between snRNA-seq and the two independent single-cell ATAC-seq technologies revealed that sci-ATAC-seq chromatin accessibility profiles (Pearson’s correlation coefficient; PCC = 0.26) were comparable predictors of transcript abundances relative to 10x Genomics scATAC-seq (PCC = 0.22), although the overall correlations to RNA abundances were relatively low (**Supplementary Figure 4**). Overall, our analysis demonstrates that sci-ATAC-seq robustly recapitulates known cell-type-specific chromatin and gene expression profiles.

### Regulatory consequences of lineage-specific chromatin accessibility

A major advantage of single-cell methods over bulk profiling is the ability to enrich signals that originate from rare cell types. To empirically test this, we identified ACRs at the bulk- and cell-type scales using sci-ATAC-seq, 10X Genomics scATAC-seq, and their union. Identification of ACRs using data partitioned by cell type resulted in 11,850 more ACRs than bulk on average, showcasing a 32% increase in sensitivity (**Figure 5A**). As a specific example, we selected a random ACR on chromosome 1 (Chr1:25450953-25451274) detected following cell-type partitioning that was undetected in the bulk sci-ATAC-seq data set (**Figure 5B**). This ACR was highly enriched (250-fold) in the phloem (accessible in 5% of phloem cells) despite being accessible in less than 0.02% of total nuclei, suggesting that a substantial portion of cell-type-specific ACRs are masked by bulk approaches (**Figure 5B**). In addition, we observed that ACRs identified following cell-type partitioning were more concordant across data sets compared to bulk-scale profiling (**Figure 5C**). Thus, interrogation of chromatin accessibility from cell-type resolved data significantly improves the sensitivity for identification of rare and cell type-specific CREs.

**Figure 5.**
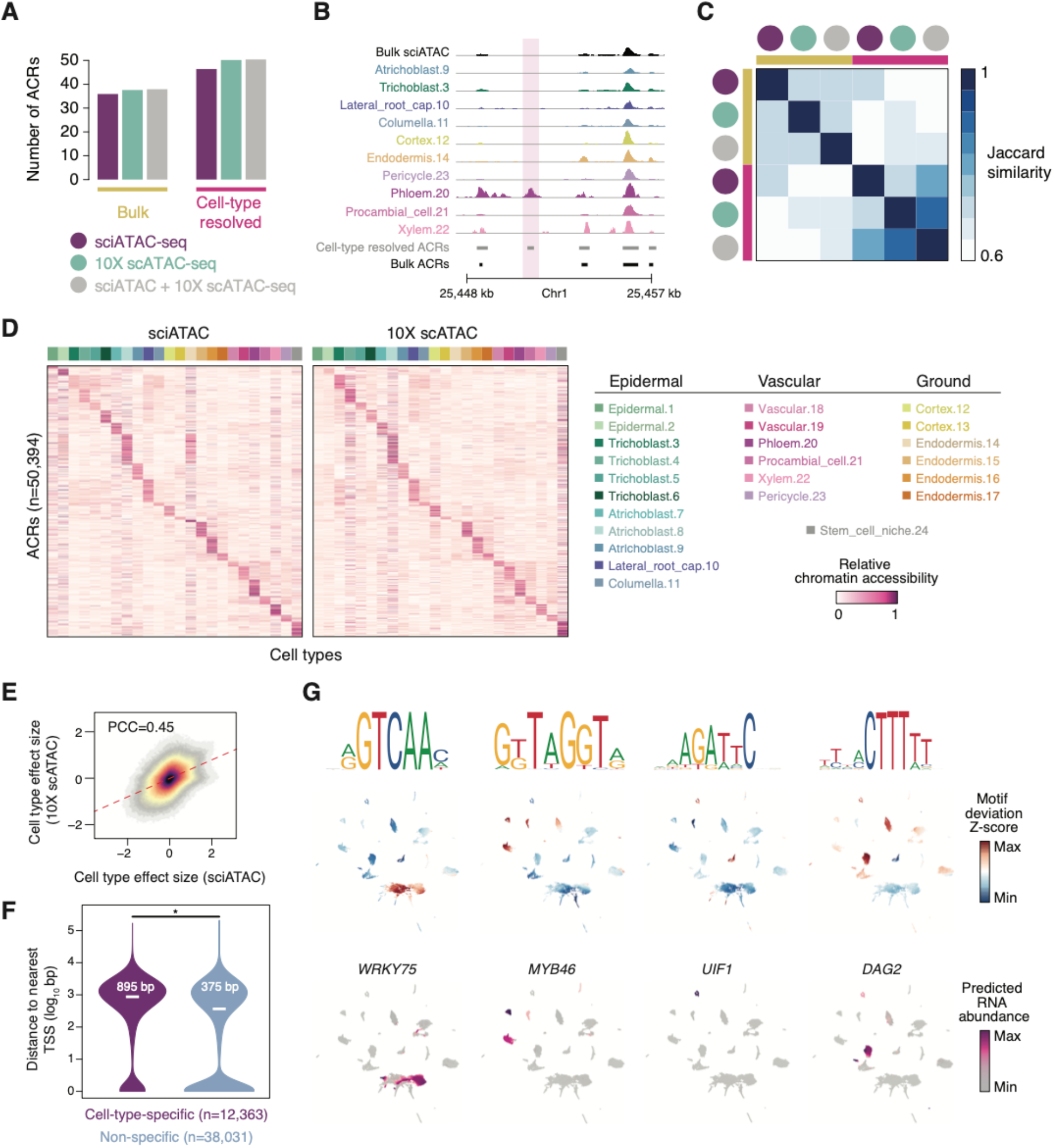
Cell type-specific ACRs and TF regulatory signatures. **(A)** The number of ACRs identified in bulk (all nuclei) compared to ACR counts identified after merging ACRs from distinct cell types for sci-ATAC-seq, 10x Genomics scATAC-seq, and sci-ATAC-seq + 10x Genomics scATAC-seq data sets. **(B)** Whole root bulk and pseudobulk cell type Tn5 integration site coverages from sci-ATAC-seq. Track heights denote read coverage scaled per million. The bottom two tracks represent ACRs from cell-type and whole root bulk resolutions, respectively. **(C)** Pairwise Jaccard similarity (fraction of overlapping ACRs versus the union of ACR sets) for ACR calls among various scATAC-seq technologies and resolutions. **(D)** Relative chromatin accessibility across sci-ATAC-seq and 10x Genomics scATAC-seq based cell types and 50,394 ACRs. **(E)** Comparison of beta values from likelihood ratio tests of the contribution of cell-type membership towards ACR chromatin accessibility status between sci-ATAC-seq and 10x Genomics scATAC-seq. **(F)** Distributions of ACR distance to the nearest TSS for cell type-specific and non-specific ACRs. **(G)** Comparison of matched motif deviation and imputed TF RNA abundance levels for four TFs with cell type-specific gene expression patterns.

To gain insight into cell type-specific *cis*-regulation, we first evaluated global patterns of chromatin accessibility among cell types for sci-ATAC-seq and 10X scATAC-seq technologies. Expectedly, distributions of chromatin accessibility across cell types were highly concordant between two different single-cell ATAC-seq technologies (**Figure 5D**). To quantify potential differences, we performed differential accessibility analysis for sci-ATAC-seq and 10X scATAC-seq, independently. The effect of cell-type on ACR accessibility status was well-correlated between technologies, indicating that the magnitude and direction of differential chromatin accessibility across cell types were consistent between sci-ATAC-seq and 10X scATAC-seq (**Figure 5E**). By applying uniform thresholds (FDR < 0.1 and Beta coefficient > 1) for each technology, we identified significant changes for ∼25% (12,363/50,394) of ACRs. Interestingly, ∼46% (5,498/11,850) of cell-type-specific ACRs were not identified within the bulk ACR set, further showcasing the usefulness of single-cell analysis for finding ACR enriched in rare cell-types. In addition, ACRs with cell-type specificity were on average nearly 2.5 times (895 bp versus 375 bp) further away from the nearest TSS compared to non-specific ACRs, suggesting that even in the small *A. thaliana* genome, distal CREs could be important contributors towards cell identity (**Figure 5F**). In summary, our approach enabled the unbiased identification of cell type-specific chromatin accessibility, is well supported by previous data, and provides a general resource for defining cell-type-specific *cis-*regulatory elements in the effort to better understand fine-scale biological processes in plants.

A longstanding question in developmental biology is defining the sets of TFs that are involved in generating and maintaining cell type diversity from an invariant genome. To further understand the transcriptional regulatory sequences underlying accessible chromatin in distinct cell identities, we performed global motif enrichment analysis across all accessible chromatin regions specific to each cell. Enrichment of motif sequences in accessible chromatin regions relative to background (motif deviation scores) for known regulators were consistent with the predicted cell identities, including *WRKY* family TFs in the specification of root epidermal progenitors (Marand et al., 2021), *MYB* in endodermis, cortex and pericycle (Cohen et al., 2020), *ULT1 interacting factor* (*UIF*) in the stem cell niche and phloem cells (Moreau et al., 2016), and *DOF AFFECTING GERMINATION* (*DAG*) in the vascular system (Gualberti et al., 2002) (**Figure 5G**). This analysis highlights global capture of accessible sequence motifs targeted by key lineage-specific TFs that are important regulators of cell identity. Furthermore, patterns of motif accessibility were supported by transcription of the cognate TF (**Figure 5G**). Gain of motif accessibility in non-expressing cells, such as *DAG2* motif accessibility in pericycle nuclei, may reflect non-cell autonomous activity as has been previously reported (Marand et al., 2021). These data showcase the ability to uncover TF-target sequences associated with lineage-specification that are important for plant growth and development.

## DISCUSSION

Cellular heterogeneity poses a major obstacle in the dissection of gene regulatory programs driving plant development. Although great progress has been made with single-cell-based analysis of chromatin status in plants by using commercial platforms, there are trade-offs between data quality, throughput and cost. Our study expands the potential application of combinatorial indexing in plant single-cell sequencing and demonstrates its usefulness to profile single-cell chromatin accessibility in *A. thaliana* root tissue. Compared with previous 10X Genomics-based data, our sci-ATAC-seq strategy increases the yield and efficiency, doesn’t require cell sorting or commercial reagents and reduces the cost to ∼$0.01 per nucleus. A relaxed sci-ATAC-seq workflow for 10-20 plates can be completed in a week to obtain chromatin accessibility information from 15,000-30,000 nuclei. The ability to safely freeze and store plates during the protocol makes this method highly modular and improves scalability.

Our analysis demonstrated that sci-ATAC-seq maps provide information-rich measurements for nuclei, enabling the identification and annotation of cell types and their underlying regulatory elements. Moreover, sci-ATAC-seq is able to reveal key gene regulatory features, such as genomic regions with cell type-specific chromatin accessibility and transcription factor motifs. In addition, the combinatorial indexing method has the potential to increase throughput to millions of cells by coupling to a droplet-based microfluidic platform for PCR, instead of using 96-well plates. In animal researches, such an approach has already been further modified to joint profile transcription or epigenomic features such as transcription factor binding and histone modification (Cao et al., 2018; Luo et al., 2017; Zhu et al., 2021). We hope that by removing cost, instrument, and other technical obstacles, sci-ATAC-seq can be readily adopted by more plant research laboratories and facilitate new insight into the complex molecular mechanisms underlying cell-specific gene expression, cell fate determination and plant growth and development.

## METHODS

### Single-cell combinatorial index ATAC-seq

A detailed step-by-step sci-ATAC-seq protocol with reagent and equipment lists was included in the supplementary information. The Tn5 expression and purification were performed as previously described (Tu et al., 2020). All plasmids could be obtained from Addgene (accession #127916). All computer codes have been deposited on GitHub.

### Nuclei isolation

Seeds of *A. thaliana* were surface sterilized and then sown on half-strength Murashige and Skoog (MS) medium plates containing 1% agar without added sucrose. Plates were vertically placed in a 22°C plant growth chamber with a photoperiod of 18 hours of light, 6 hours of dark for two weeks. The seedlings were grown for two weeks before nuclei isolation. One of the key technical challenges for scATAC-seq with plant tissue is the clumping of broken nuclei. As detergents, such as Triton X-100 and Tween 20, that are used to lyse plant organelles, also disrupt the outer nuclear membrane. We previously observed that plant nuclei after formaldehyde fixation are highly resistant to clumping, and remain intact after heat shock, SDS denaturation, centrifugation and over-night incubation steps during *in situ* Hi-C library preparation (Dong et al., 2017). Since it has been reported that ATAC-seq can be performed on formaldehyde fixed animal tissues (Chen et al., 2016), we introduced a short fixation to the nuclei isolation process before lysing the plant organelles with detergent to preserve the nuclear membrane and prevent clumping during the single-cell library preparation. The ability to use detergent washes to lyse organelle without affecting nuclear integrity enabled us to reduce doublet rates and remove organelle DNA contamination without the need for cytometrical sorting.

To isolate intact nuclei, roots from ∼40 Arabidopsis seedlings were first immersed in 1X PBS with 1% formaldehyde and 1.5 mM PMSF. The solution was vacuumed twice for 5 min, and the roots were washed with distilled water. The fixed tissues were then transferred into a Petri dish with 10 mL pre-chilled Cutting Buffer and chopped into small pieces to release the nuclei. The solution was then filtered through a 40 μm cell strainer and nuclei were collected by centrifugation at 500g, 4°C for 15 min. The pellet was then resuspended with Cutting Buffer without formaldehyde and filtered through a 10 μm nylon filter. The nuclei were then pelleted again by centrifugation and washed one more time with the Cutting Buffer. Before the final centrifugation, a small fraction of the solution was removed and stained with SYBR Green to determine nuclei concentration. We recommend using 2-5 × 10^5^ nuclei for each sci-ATAC-seq experiment as using too many nuclei will increase clumping during the centrifugation steps. The nuclei pellet was then washed once with 5 mL of Tagmentation Buffer without DMF, and resuspended in 5 mL of Tagmentation Buffer before adding to the 4 PCR plates with unique indexed Tn5 in each well.

### Tn5 tagmentation and split-pool-split

The sci-ATAC-seq depends on the Tn5 transposon to introduce barcoded adapters into individual nucleus for subsequent demultiplexing the single cell data from the bulk sequencing reads. For such transposon-based ATAC-seq, it is often necessary to experimentally determine an optimized Tn5 concentration to avoid over or under tagmentation (Orchard et al., 2020). This is particularly challenging for sci-ATAC-seq, as nuclei would have to be tagged by hundreds of differentially barcoded transposons. We used the hyper-stable TS-Tn5 transposase that fused Tn5 to the *E. coli* chaperone elongation factor (EF-Ts) (Tu et al., 2020). As it has been shown that TS-Tn5 could limit transposition into inaccessible chromatin due to the increased protein size, simultaneously simplifying the assay and improving the signal-to-noise ratio. We observed that read pileup around genes from our method appeared more discrete than those from 10X Genomics scATAC-seq, consistent with the hypothesized reduction in background integrations afforded by the TS-Tn5 tagmentation enzyme (Figure 2 F-G; Supplementary Figure 2). To obtain a combination of 384 indexed transposon, 24 forward Tn5-ME-A and 16 reverse Tn5-ME-B adapters were used to assemble the transposon at 25°C for 1 hour. First, 1.5 μL of each A and B transposons were distributed to four 96-well plates (each well contains a unique combination of A and B indexed Tn5). The nuclei in 12 μL Tagmentation Buffer were added to each well of the four 96-well plates. The plates were then sealed and the tagmentation reaction was carried out for 45-60 minutes at 37°C with occasional shaking. The reaction was stopped by flushing each well with 100 μL Wash Buffer supplemented with EDTA to quench the Mg^2+^. All the nuclei were transferred to a reservoir and to a 14 mL round-bottom polystyrene tube. It was then pelleted and washed once with the Wash Buffer. The nuclei pellet was then resuspended to a concentration of ∼10 nuclei per microliter. 1.5 μL of nuclei solution was then pipetted into each well of 96-well plates. 3 μL of Lysis Buffer containing SDS was then added to each well to lyse the nuclei. The plates were then sealed and could be stored at -20 °C.

### Library amplification

To perform plate PCR with unique row and column barcode primers, the stored nuclei plates were briefly centrifuged and heated to 50 °C in a PCR machine for 5 min before adding 7.5 μL Row PCR Master Mix containing 2% Tween-20 and 7.5 μL Column PCR Master Mix containing DNA polymerase to reconstitute a 20 μL PCR reaction in each well. The detergent in the Row PCR Master Mix is required to quench the SDS in the Lysis buffer. The plate was then sealed and the tagged DNA was PCR amplified to acquire the second barcode that labeled DNA from each well. The 96 PCR reactions were pooled and purified using a Qiagen MinElute Kit on a vacuum manifold. The purified PCR products were re-amplified using modified Illumina TruSeq PCR primers (Supplementary Method and Tables), which gave the tagged DNA a third barcode to label each plate. Libraries prepared from 10 plates (5 for each biological replicates) were sent for Illumina HiSeq × sequencing using the manufacturer’s conventional sequencing reagent and primers. It should be noted that if HiSeq × is used for sequencing, 5-10% spike-in library (e.g. PhiX control from Illumina) must be added to the lane to balance the nucleotide distribution at the beginning of the forward and reversed reads. Spike-in control is not required when the sci-ATAC-seq libraries are sequenced on the latest NOVAseq system, where different forms of libraries are mixed and sequenced in one lane.

### Data processing

Cutadapt (version 2.10) was used to demultiplex the inline barcode of the sci-ATAC-seq. The 10x Genomics scATAC-seq data was processed as previously described (Marand et al., 2021). Barcode index sequences were appended to the read names of paired-end reads using the extract command of UMItools v1.0.1 with non-default parameters (--bc-pattern=NNNNNNNNNNNNNNNNNNNNNNNNNNN) and aligned to the *A. thaliana* TAIR10 reference genome with BWA mem v0.7.17. Aligned reads were filtered with samtools v1.6.0 to remove low quality alignments (samtools view -q 30), non-properly paired reads (samtools view -f 3), duplicate reads with the picard tools v2.21.6 function MarkDuplicates and multiple mapped reads (any read pair associated with the XA tag appended by BWA mem) with a simple PERL script. The remaining alignments were converted to single-base resolution Tn5 integration sites by adjusting the start position of forward and reverse strand alignments by +4 and -5, respectively. Only unique Tn5 insertion sites per barcode were retained.

### Nuclei identification and quality control

Nuclei identification and quality control steps were implemented with the R package, *Socrates* v0.0.1. Briefly, single-base resolution Tn5 insertion sites for each data set were loaded in *Socrates* with TAIR10 GFF gene annotation and TAIR10 genome index. Bulk-scale ACRs were identified within Socrates using callACRs (genomesize=9.5e7, shift= -50, extsize=100, fdr=0.05) for estimating fraction of Tn5 integration sites within ACRs for each nucleus. Meta data for each nucleus was collected using the buildMetaData function with a TSS window size of 2-kb (tss.window=2000). Thresholds for the minimum number of unique Tn5 insertion sites, fraction of Tn5 insertion sites within 2-kb of TSSs and fraction of Tn5 insertion sites within ACRs were identified manually for each independent data set (minimum of 1,000 Tn5 insertion sites per nucleus, 70% of Tn5 insertions within 2-kb and 50% of Tn5 insertions within ACRs across all data sets). Doublet enrichment scores were estimated in *Socrates* using the approach previously described (Granja et al., 2021).

### snRNA-seq data processing

Raw snRNA-seq data from *A. thaliana* roots were acquired from NCBI GEO GSE155304. Raw reads were aligned to the TAIR10 reference genome and processed with cellranger v4.0.0. snRNA-seq quality control and nuclei identification were performed as previously described (Marand et al., 2021).

### snRNA-seq, scATAC-seq, and sci-ATAC-seq data integration

UMI and Tn5 insertion site counts matrices were generated from gene body and gene body + 500bp upstream TSSs for snRNA-seq and single-cell ATAC-seq data sets, respectively, and loaded into R using liger. Each data set was normalized independently using the function normalize. Highly variable genes were selected based on the substructure observed in snRNA-seq with the function selectGenes (datasets.use=“snRNA”) and used to subset and scale all data sets via the function scaleNotCenter. A joint embedding containing snRNA-seq, scATAC-seq, and sci-ATAC-seq nuclei was generated using integrative non-negative matrix factorization (iNMF) with the function optimizeALS (k=15, lambda=5). The metagene factors estimated by iNMF were then quantile normalized with the function quantile_norm (do.center=F, ref_dataset=“snRNA”) using the snRNA-seq nuclei as the reference to fully integrate all data sets into a shared embedding. Groups of nuclei with similar patterns of gene expression and chromatin accessibility were identified with the function louvainCluster with non-default parameters (resolution=0.25, k=35, eps=0, prune=1/10). Nuclei-nuclei relationships were visualized by reducing the dimensions of the iNMF loadings with runUMAP (n_neighbors=35, min_dist=0.01). RNA abundances were predicted for each scATAC-seq and sci-ATAC-seq nucleus by averaging the expression levels of the 20 nearest snRNA-seq nuclei neighbors using the function.

### Independent clustering analysis for sci-ATAC-seq and 10X scATAC-seq

Independent analysis of sci-ATAC-seq and 10x scATAC-seq data was performed using the R package, *Socrates* (Marand et al. 2021, *Cell*). Briefly, a sparse nuclei by 500-bp window matrix scored by presence/absence of Tn5 insertions was filtered to remove nuclei with less than 100 accessible windows and windows with less than 0.1% of nuclei with a Tn5 insertion. The filtered matrices were normalized using a quasibinomial logistic regression regularized model with the default settings of the function *regModel*. The dimensionality of the normalized accessibility scores were reduced using the function *reduceDims* (n.pcs=50, cor.max=0.8). The reduced embedding was visualized as a UMAP embedding using *projectUMAP* (k.near=35, m.dist=0.05). Finally, we identified graph-based clusters using the Louvain neighborhood clustering algorithm with *callClusters* (res=0.6, threshold=5, e.thresh=3, k.near=35). Due to lab-specific batch effects in the 10X scATAC-seq data, we used the R package, *Harmony* (Korsunsky et al. 2019. *Nature Methods*) to remove batch effects using a covariate for lab of origin prior to graph-based clustering with non-default parameters (do_pca=F, max.iter.harmony=30, theta=2, lambda=0.1). Adjusted Rand Index scores were determined using the R package, *mclust*. Reciprocal NNLS between sci-ATAC-seq clusters and 10x scATAC-seq clusters was performed as previously described (Domcke et al. 2020. *Science*) using the R package, *nnls*.

### Cell-type annotation

To enable cell-type annotations for each individual nucleus, we devised an annotation strategy that estimates the relative enrichment of cell-type-specific markers against permuted sets of randomly selected cell-type-specific genes. Specifically, we collected a list of known cell-type-specific marker genes for various cell types present in the root of *A. thaliana* and estimate Z-scores of gene expression for these genes across nuclei specifically using the snRNA-seq data set. For each cell-type, we estimated the average Z-score from × markers specific to the given cell type. Then, we derived 1,000 random sets of × cell-type-specific markers, excluding the cell type of interest, estimating the average Z-score of each randomized set. The relative cell-type-specific enrichment was then estimated as follows (Eq. 1):

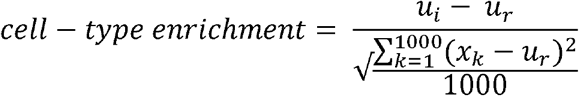

Where *i* denotes the average Z-score for the cell type of interest, r is the average Z-score from the randomized permuted gene sets and k is the index for the vector of permuted gene sets. Owing to the high degree of sparsity and drop-outs in single-cell experiments, we smoothed the cell-type enrichment scores using a diffusion-based weighted affinity matrix derived from the iNMF loadings, 25 nearest neighbors, and three diffusion time steps using a previously described approach for imputing single-cell gene expression values (citation). Briefly, we first computed a nucleus-nucleus distance matrix using the iNMF loadings and transformed distances into nucleus-nucleus affinities with a Gaussian kernel function. The affinity matrix was then symmetrized to reduce noise and the effect of outliers by addition of the transposed affinity matrix and rescaling the rows to sum to one, resulting in a Markov transition matrix (Mi,j) that represents the probability of a transition between nucleus i and j. To remove sources of technical noise, including dropouts and under-sequencing, we use “diffusion time”, t, to raise M to the power t. Smoothed cell-type enrichment scores were estimated by matrix multiplication of the Markov transition matrix with the original nucleus by cell type matrix. The cell-type enrichment scores were converted into a probability distribution by rescaling by the sum across all possible cell types. Nuclei were labeled with the cell-type with the greatest probability.

To extend snRNA-seq cell type annotations to scATAC-seq and sci-ATAC-seq within the same embedding, we trained a weighted knn model on the snRNA-seq cell type annotations using the iNMF loadings using the R function kknn and assigned cell type annotations based on the highest probability. From the full embedding, clusters were labeled as the cell type with the greatest frequency among nuclei. Automated cell-type annotations were assessed and revised as necessary by computing differential gene expression and accessibility and visualizing normalized gene expression and accessibility values on the reduced UMAP embedding as previously described.

### ACR identification

ACRs were first identified using the pseudobulk Tn5 integration sites (ignoring barcode information) for droplet-based scATAC-seq and sci-ATAC-seq independently with MACS2 v2.2.7.1 (--nomodel --keep-dup all --extsize 100 --shift -50 --qvalue 0.05). ACRs for scATAC-seq and sci-ATAC-seq were then filtered using an empirical FDR method (FDR < 0.01) as previously described to (Hufford et al. 2021. Science). Following nuclei clustering and annotation, we performed an additional round of ACR identification (using the same parameters and filtering steps as described above) for each cluster using all Tn5 integrations sites regardless of technology (scATAC-seq + sci-ATAC-seq, hereafter referred to as “combined scATAC-seq”) as well as single-cell ATAC-seq technology in isolation by aggregating Tn5 integration sites for barcodes associated with the focal cluster, generating ACR calls from each cluster for the combined scATAC-seq, and scATAC-seq and sci-ATAC-seq in isolation. Cluster-based ACRs were merged into single BED files for the combined data set, scATAC-seq and sci-ATAC-seq, independently, using the BEDtools merge command. Concordance among bulk, cell-type resolved, and single-cell ATAC-seq technologies was performed using the BEDtools jaccard function. Differential chromatin accessibility was performed as previously described with the exception that all nuclei were included in the per ACR and per cluster model (replacing the randomly selected nuclei as a reference set). In addition, the differential accessibility logistic regression models were determined for scATAC-seq and sci-ATAC-seq nuclei, independently. The beta values (effect sizes) from the likelihood ratio tests (between a model including cell type membership and nuclei read depth and a model with only nuclei read depth) were extracted for each ACR and cell type for scATAC-seq and sci-ATAC-seq data sets, respectively. Thus, the effect sizes represent the contribution of cell-type identity towards the relative chromatin accessibility of a locus conditioned by the cognate single-cell ATAC-seq technology. Finally, we defined differential accessibility as ACRs with FDR < 0.1 and beta values > 1 in both data sets, resulting in a total of 12,363 differentially accessible chromatin regions and an average of 515 per cell type.

### Transcription factor motif analysis

To determine the relative enrichment of various TF motifs within accessible chromatin regions for each nucleus, we first constructed a sparse binarized matrix of ACRs by nuclei by scoring the presence/absence of Tn5 integration sites for each nucleus using the combined scATAC-seq ACRs with differential chromatin accessibility (FDR < 0.05 and log2FC > 2). The sparse ACR by nuclei matrix was filtered to retain ACRs that were accessible in at least 100 nuclei followed by removing nuclei with less than 100 accessible ACRs and a fraction of Tn5 insertions within the remaining ACRs less than 0.1. JASPAR 2020 motif occurrences within differentially accessible ACRs were collected using the function matchMotifs from the R package chromVAR based on significant motif matches (default P-value = 5e-5) to the underlying ACR sequences. Motif deviation scores for each nucleus were then estimated using the deviationScores function after normalizing by GC bias. To visualize motif deviations and remove the effects of variable sequencing coverage and point density on the UMAP embedding, motif deviations were smoothed using the diffusion-based affinity matrix constructed from the multiple modality integrated iNMF loadings and subsequently transformed to Z-scores.

## ACCESSION NUMBERS

The sequencing data from this data have been deposited in the NCBI SRA under the accession codes PRJNA758591. The R source code and package used throughout the analysis can be found available in the following GitHub repository: https://github.com/plantformatics/plant_sciATAC.

## AUTHOR CONTRIBUTIONS

R.S. and S.Z. designed and supervised the project; X.T. and S.Z. conducted the experiments. A.P.M. performed computational analysis. X.T. and A.P.M. wrote the manuscript. All the authors reviewed and contributed to the paper.

## ACKNOWLEDGMENTS

R.J.S. is a co-founder of REquest Genomics, a company that provides epigenomic services. The remaining authors declare no competing interests.

## Supplementary information

Supplementary Method

Supplementary Figures 1-3

Supplementary Tables 1-2

