## Supplementry information for "A combinatorial indexing strategy for epigenomic profiling of plant single cells"

**The PDF file includes:**

Supplementary Methods

Supplementary Figures 1-4

Supplementary Tables 1-3

### Supplementary Methods

I. **Adapter and primer design:**
1) Tn5-ME-A index adapter A1-A24:

```
5’-GGCGGTAGGCGTG NNNNN AGATGTGTATAAGAGACAG-3’
     3’-TCTACACATATTCTCTGTC-phos-5’
```

2) Tn5-ME-B index adapter B1-B16

```
5’-CCAACACCCGTGCG NNNNN AGATGTGTATAAGAGACAG-3’
                     3’-TCTACACATATTCTCTGTC-phos-5’
```

3) Plate PCR row index primer A-H

5’-ACACTCTTTCCCTACACGACGCTCTTCCGATCT NNNNNN GGCGGTAGGCGTG-3’

4) Plate PCR column index primer 1-12

5’-GTGACTGGAGTTCAGACGTGTGCTCTTCCGATCT NNNNNN CCAACACCCGTGCG-3’

5) Illumina TruSeq PCR Forward primer (N50x index)

5’-AATGATACGGCGACCACCGAGATCTACAC NNNNNNNN ACACTCTTTCCCTACACGAC GCTCTTCCGATCT-3’

6) Original Illumina TruSeq PCR reverse primer (PCR2.x index)

5’-CAAGCAGAAGACGGCATACGAGAT NNNNNN GTGACTGGAGTTCAGACGTGTGCTC TTCCGATCT-3’

7) New Illumina TruSeq PCR reverse primer (N70x index)

5’-CAAGCAGAAGACGGCATACGAGAT NNNNNNNN GTGACTGGAGTTCAGACGTGTGCTC TTCCGATCT-3’

The barcode position is indicated by the N colored in red. The oligo sequences are shown in Supplementary Table 3. Note that our libraries were prepared using the older version of Illumina TruSeq PCR reverse primer with 6 nt PCR2.x index sequence, while the latest version of the Illumina N70x index has 8 nt barcode. Both can be sequenced in HiSeq and NOVAseq machines. But we would suggest new users to use the latest N70x index to take full advantage of the longer index read length of the new Illumina machines.

**II. Anneal index Tn5 adapters**

1. Dissolve 24 Tn5-ME-Ax, 16 Tn5-ME-Bx and the complimentary Tn5-ME-Rev single stranded DNA oligonucleotides (oligos) with TE to 100 μM.
2. In three 8-well-PCR strips, mix equal volume (5 μL) of Tn5-ME-Ax and Tn5-ME-Rev in a PCR tube to obtain a 50 μM adapter stock.
3. In two 8-well-PCR strips, mix equal volume of Tn5-ME-Bx and Tn5-ME-Rev as above.
4. Heat the to 95°C in a PCR machine and gradually lower the temperate to 25°C by cycling (-1°C per 10 sec cycle).
5. Add TE (70 μL) to each well to dilute the 50 μM adapters to 6.25 μM, and store the 24 Tn5-ME-A and 16 Tn5-ME-B adapter in PCR strips prior to use of the protocol (Note: we often store them as aliquot for one Tn5 assembly).

**III. Assemble 384 index TS-Tn5**

1. The TS-Tn5 expression and purification were performed as previously described (Tu et al., 2020). All plasmids could be obtained from Addgene (accession #127916).
2. Prepare 2 mL **Tn5 Dilution buffer** in a 2 mL tube:

| Reagent | Amount for 2 mL | Final Concentration |
| --- | --- | --- |
| Glycerol | 1.26 g | 50% |
| NaCl (4 M) | 150 μL | 0.3 M |
| Tris (1M pH 8.0) | 10 μL | 5 mM |
| EDTA (0.5 M) | 0.4 μL | 0.1 mM |
| DTT (100 mM) | 10 μL | 1 mM |
| Water | 815 μL |  |

3. Dilute TS-Tn5 to 0.5 OD^280^ with **Tn5 Dilution Buffer**. Prepare 180 μL diluted Tn5.
4. To assemble TS-Tn5 with A1-A24 adapters, mix 3.6 μL Ax adapter (6.25 μM) with 3.6 μL diluted TS-Tn5 in three 8-well-PCR tube strips.
5. To assemble Tn5 with B1-B16 adapters, mix 5.2 μL Bx adapter (6.25 μM) with 5.2 μL diluted Tn5 in two 8-well-PCR tube strips.
6. Incubate the PCR strips at 25 °C for 1 h to assemble the transposon.
7. Add 19.8 μL **Tn5 Dilution Buffer** to the assembled Tn5 with adapter A, and mix well.
8. Add 28.6 μL **Tn5 Dilution Buffer** to Tn5 with adapter B, and mix well.
9. Distribute 1.5 μL TS-Tn5 with adapter B1-16 to each column of the four PCR 96 well reaction plates on ice. The volume of the assembled Tn5 is 39 μL that is sufficient for 24 wells with 2 to spare.
10. Distribute 1.5 μL TS-Tn5 with adapter A1-24 to each row of the four PCR 96 well reaction plates on ice. The volume of assembled Tn5 is 27 μL that is sufficient for 16+2 wells.
11. After adding Tn5 with adapter Ax and Bx, each well now contain 3 μL of assembled Tn5. The four plates can be used immediately or sealed for overnight storage at -20 °C.

**V. Prepare Nuclei and ATAC-seq**

1. Roots from ~40 Arabidopsis seedlings were harvested and immersed in 1X PBS with 1% formaldehyde and 1 mM PMSF. The solution was then vacuumed infiltrated twice for 5 min, and the roots were washed with distilled water.
2. Transfer the fixed root tissues to Petri dishes with 10 mL chilled **Cutting Buffer** and cut into thin slices (usually less than 1 mm thick) under the fume hood.

**Recipe for 40 mL Cutting Buffer (kept on ice):**

| Reagent | Amount for 40 mL | Final Concentration |
| --- | --- | --- |
| Tris (1M pH 7.5) | 400 μL | 10 mM |
| NaCl (4 M) | 100 μL | 10 mM |
| MgCl_2_ (1 M) | 120 μL | 3 mM |
| CaCl_2_ (1 M) | 40 μL | 1 mM |
| Hexylene Glycol | 40 μL | 0.1% |
| BSA (10%) | 400 uL | 0.1% |
| PVP-40 (10%) | 40 μL | 0.01% |
| NP-40 (10%) | 40 μL | 0.01% |
| Spermidine (0.5 M) | 40 μL | 0.5 mM |
| Spermine (0.2 M) | 40 μL | 0.2 mM |
| PMSF (0.2 M) | 200 μL | 1 mM |
| β-mercaptoethanol | 40 μL | 0.1% |
| Water | Top up to 40 mL |  |

3. Filter through a 40 μm cell strainer, transfer to a 14 mL Falcon round-bottom polypropylene tube on ice.
4. Note that for soft tissues such as Arabidopsis roots that are difficult to cut, a pre-fixation in 1% formaldehyde under vacuum for 10 to 15 min will harden the tissues and make them easier to process. For waxy tissues such as maize and rice leaves that are difficult to fix, it is best to cut the unfixed tissues. We would add 1.25 mL 4% formaldehyde to the filtered crude nuclei suspension to a final concentration of 0.5%, mix well and incubate on ice for 5 min. Then add 0.5 mL Tris buffer (1 M, pH 7.5) and 0.5 mL 2 M Glycine to quench the formaldehyde.
5. Add 0.1 mL of 20% Triton X-100 (prepared monthly and stored in 4°C fridge) to a final concentration of 0.2% to lyse the chloroplasts and mitochondria, mix well and stand on ice for 2 min.
6. Remove 1 uL of the solution and stain the nuclei with SYBR-green to check the nuclei quality and concentration under a microscope. Insufficient fixation or using too many nuclei (more than 1x10^6^) would cause clumping. We normally aim for 1 to 5x10^5^ nuclei for each preparation.
7. Pellet the nuclei by centrifugation (swinging-bucket centrifuge rotor) at 500 x g for 15 min at 4°C.
8. Remove the supernatant using a dropper. To avoid disturbing the nuclei, we often leave ~0.5 mL supernatant on top of the pellet.
9. Resuspend the nuclei pellet gently with 1 mL **Cutting Buffer**, filter through a 10 μm pore size Nylon membrane filter. Flush the filter with 3 mL of **Cutting Buffer** (Note that one should choose filter with different pore size according to the size of the nuclei).
10. Transfer all to a new 14 mL Falcon tube and spin down at 500 x g for 10 min at 4°C.
11. Resuspend the nuclei pellet gently with 5 mL **Cutting Buffer** and spin down at 500 x g for 10 min at 4°C
12. Resuspend the nuclei pellet with 5 mL chilled 1X **Tagmentation Buffer** without DMF and supplemented with 0.1% BSA and 0.01% NP-40. Spin down at 500 x g for 10 min at 4°C.

**Recipe for 5 mL Tagmentation buffer w/o DMF (kept on ice):**

| Reagent | Amount for 5 mL | Final Concentration |
| --- | --- | --- |
| Tris (1M pH 7.8) | 100 μL | 20 mM |
| KAc (1 M) | 300 μL | 60 mM |
| MgCl_2_ (1 M) | 50 μL | 10 mM |
| BSA (10%) | 50 uL | 0.1% |
| NP-40 (10%) | 5 μL | 0.01% |
| Water | 4.5 mL |  |

13. Resuspend nuclei with 5 mL 1X **Tagmentation Buffer** with 15% Dimethylformamide (DMF) and 1 x Pierce Protease Inhibitor without EDTA (PI, Invitrogen A32965).
14. Transfer the nuclei to 12 PCR tubes and use a 12-channel pipette to distribute 12 μL nuclei to each well of the four 96-well-PCR plates with assembled Tn5 transposome. Mix well by pipetting.
15. Incubate the four plates at 37°C for 45-60 min with occasional shaking.

**Recipe for 5 mL Tagmentation buffer with DMF:**

| Reagent | Amount for 5 mL | Final Concentration |
| --- | --- | --- |
| Tris (1M pH 7.8) | 100 μL | 20 mM |
| KAc (1 M) | 300 μL | 60 mM |
| MgCl_2_ (1 M) | 50 μL | 10 mM |
| DMF (100%) | 0.75 mL | 15% |
| Water | 3.8 mL |  |
| PI (100x) | 50 μL | 1x |

**VI. Pool and split nuclei**

1. Add 5 mL **Wash Buffer** to a V-bottom reservoir and add 0.4 mL extra EDTA for chelating the 10 mM Mg in the Tn5 reaction buffer.

**Recipe for 20 mL Wash Buffer (keep on ice):**

| Reagent | Amount for 20 mL | Final Concentration |
| --- | --- | --- |
| Tris (1M pH 8.0) | 100 μL | 5 mM |
| NaCl (4 M) | 100 μL | 10 mM |
| EDTA (0.5 M) | 40 μL | 1 mM |
| Hexylene Glycol | 20 μL | 0.1% |
| BSA (10%) | 200 uL | 0.1% |
| NP-40 (10%) | 20 μL | 0.01% |
| Spermidine (0.5 M) | 20 μL | 0.5 mM |
| Spermine (0.2 M) | 20 μL | 0.2 mM |
| Water | Top up to 20 mL |  |

2. Use a multichannel pipette to add 150 μL **Wash Buffer** to the four PCR plates row by row to stop the tagmentation. Transfer the nuclei back to the same reservoir each time. Optional, flush the plates again to recover all the nuclei.
3. Transfer the nuclei solution (~ 10 mL) into a new 14 mL round-bottom tube. Spin down the nuclei at 500 x g for 15 min at 4°C. Spin down the nuclei at 500 x g for 15 min at 4°C.
4. Wash the nuclei with 5 mL chilled Wash Buffer and spin down the nuclei at 500 x g for 10 min at 4°C.
5. Carefully aspirate the supernatant and resuspend in 1 mL **Wash Buffer**. Filter through a 10 μm Nylon membrane filter. Note that this filtering step could help to remove some of the clumped nuclei.
6. Remove some nuclei for SYBR-green staining and check the quality and concentration under microscope.
7. Dilute the nuclei to 10 nuclei per μL using **Wash Buffer**.
8. Use a multichannel pipette to add 1.5 μL Nuclei to each well of the PCR plates. Note that we often prepared 10 PCR plates, which is sufficient to generate 15,000 sci-ATAC-seq profiles.
9. Add 3 uL **Lysis Buffer** to each well of the PCR plate. Seal the plates, snap spin to pellet the nuclei and **Lysis Buffer** to the bottom.
10. Incubate the plates at 37°C oven for 30 min to allow the SDS to denature the Tn5 protein. The plates can then be stored at -20°C freezer. Note that we have stored them for over 6 months.

**Recipe for 10 mL Lysis buffer**

| Reagent | Amount for 10 mL | Final Concentration |
| --- | --- | --- |
| Tris (1M pH 8.0) | 50 μL | 5 mM |
| SDS (10%) | 50 μL | 0.05 % |
| Yeast carrier RNA (1 mg/mL) | 10 μL | 1 ng/uL |

**VII. PCR master mix setup for two 96-well plates**

1. Remove two nuclei plates from the storage. Span spin the plates and incubate them in PCR machines at 42 °C with heated lid for 5 min while preparing the PCR mater mixes.
2. Dispense 2.4 μL of row index primers (row A to H) (100 μM) to one 8-tube strip.
3. Aliquot 182 μL PCR master mix A into each well of the 8-tube strip:

**PCR Master Mix A for two 96-well plates**

| Reagent | Amount for 1.5 mL | Final Concentration |
| --- | --- | --- |
| Q5 Buffer (5x) | 500 μL | 1.66x |
| dNTP (10 mM) | 30 μL | 0.2 mM |
| Tween 20 (10%) | 300 μL | 2% |
| Water | 670 μL |  |

4. Dispense 1.6 μL of column index primers (column 1 to 12) (100 μM) to one and a half 8-tube strips.
5. Aliquot 122 μL PCR master mix B into each well of the 12-tube strip from above with the multiplying amount of 1.5 mL:

**PCR Master Mix B for two 96-well plates**

| Reagent | Amount for 1.5 mL | Final Concentration |
| --- | --- | --- |
| Q5 Buffer (5x) | 300 μL | 1x |
| dNTP (10 mM) | 30 μL | 0.2 mM |
| Tween 20 (10%) | 15 μL | 0.1% |
| Water | 1140 μL |  |
| Q5 hot start (2U/uL) | 15 μL | 1 U per 50 uL |

6. Use a 8-channel pipette to dispense 7.5 μL of PCR master mix A to the side of the wells without touching the nuclei lysis in each column of the two 96-well plate. Gently tap the plates to get all the liquid to the bottom. Snap spin the plates if needed.
7. Let the plates to sit at room temperature for 1 to 2 min to enable Tween-20 to quench the SDS. The plates can then be kept on ice before adding the 2^nd^ master mix.
8. Use a 12-channel pipette to dispense 7.5 μL PCR master mix B to the side of the wells.
9. Seal the plates and snap spin to get all the liquid to the bottom.
10. Transfer the PCR plates from ice to the PCR machine with the lid pre-heated to 105°C and run the thermocycling as follow:

Gap filling for 10 min at 72°C

Initial denaturation for 1 min at 80°C

Denature for 15s at 98°C

10 cycles	Anneal primers for 30s at 63°C

Extend DNA for 20s at 72°C

Final extension for 3 min at 72°C

4°C Hold

11. Add 8 mL Qiagen DNA binding buffer PB and 0.5 μL 1 μg/μL Yeast carrier tRNA to a V-shape reservoir. Transfer the PCR product (~2 mL) from one plate to the reservoir with a multichannel pipette. Flush the wells with PB to recover all the PCR products.
12. Purify the PCR product using Qiagen MinElute PCR Purification Kit on QIAvac 24 Plus vacuum manifold attached to a pump. One could also use a centrifuge machine by loading the DNA-PB solution multiple times onto the spin column.
13. Elute the DNA with 40.3 μL water and prepare the 2nd round of PCR as followed:

| Reagent | Amount for 60 uL |
| --- | --- |
| Q5 Buffer (5x) | 12 μL |
| dNTP (10 mM) | 1.2 μL |
| DNA template | 40.3 μL |
| TruSeq Forward PCR Primer N50x (10 uM) | 3 μL |
| TruSeq Reverse PCR Primer N70x (10 uM) | 3 μL |
| Q5 (2U/uL) | 0.5 uL |

14. Run the thermocycling as follow:

Initial denaturation for 1 min at 95°C

Denature for 15s at 98°C

8-10 cycles	Anneal primers for 30s at 65°C

Extend DNA for 20s at 72°C

Final extension for 3 min at 72°C

15. Purify the PCR product using 1 volume of AMPure XP beads and elute libraries with 30 uL TE. The expected yield is ~200 ng of library DNA after PCR. Optional: to avoid over amplification, we often take 10 uL of the PCR mix, add 0.25 uL EVA-green dye, and run them in a benchtop qPCR machine to determine the optimal cycle number before amplifying the remaining 50 μL PCR mix.

Video recordings showing critical steps could be viewed on: <https://space.bilibili.com/511150037/channel/detail?cid=179138>

**Supplementary Figures**

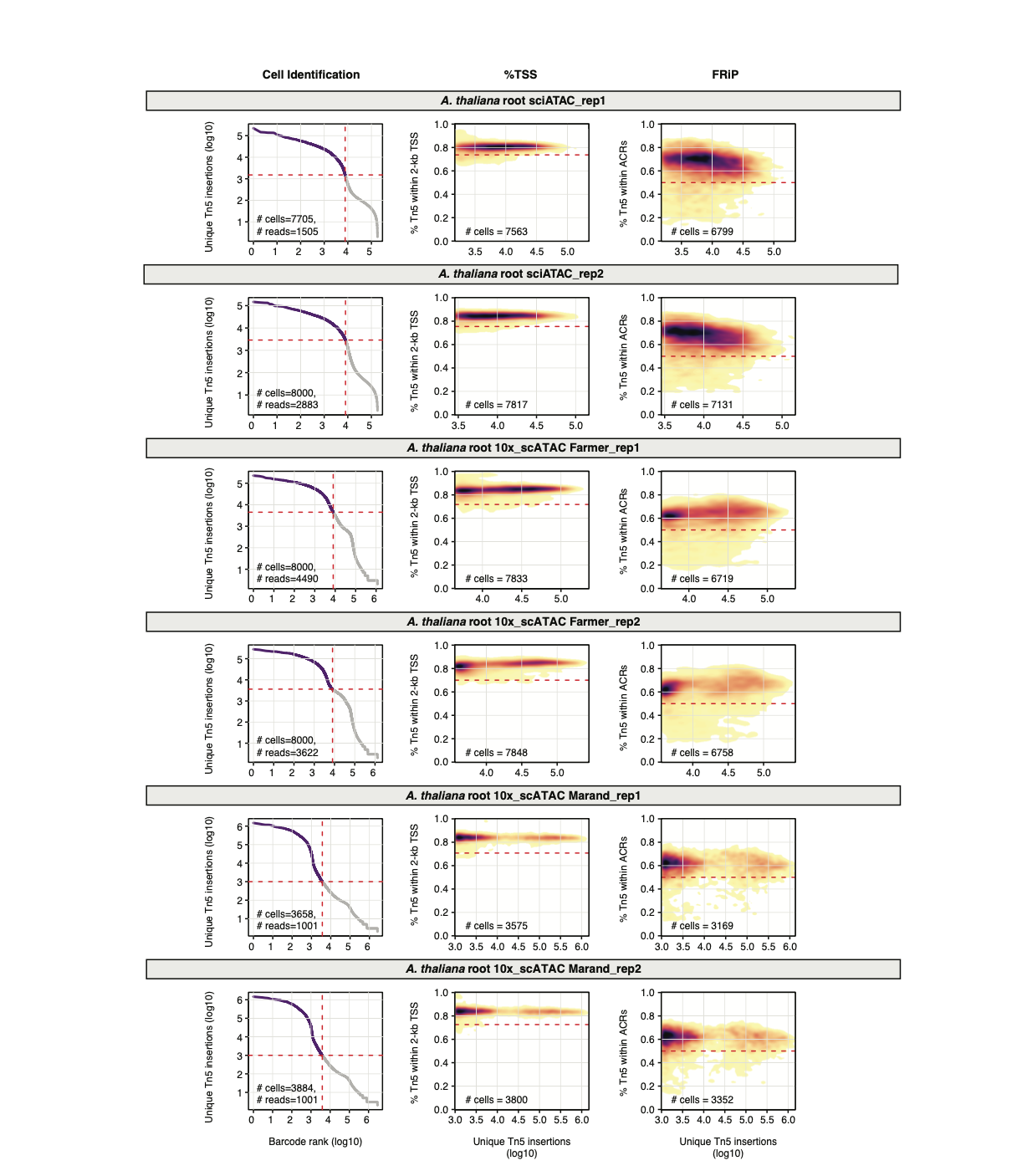

**Supplementary Figure 1.** Comparison of nuclei filtering metrics across sci-ATAC-seq and 10X Genomics scATAC-seq data sets by replicate.

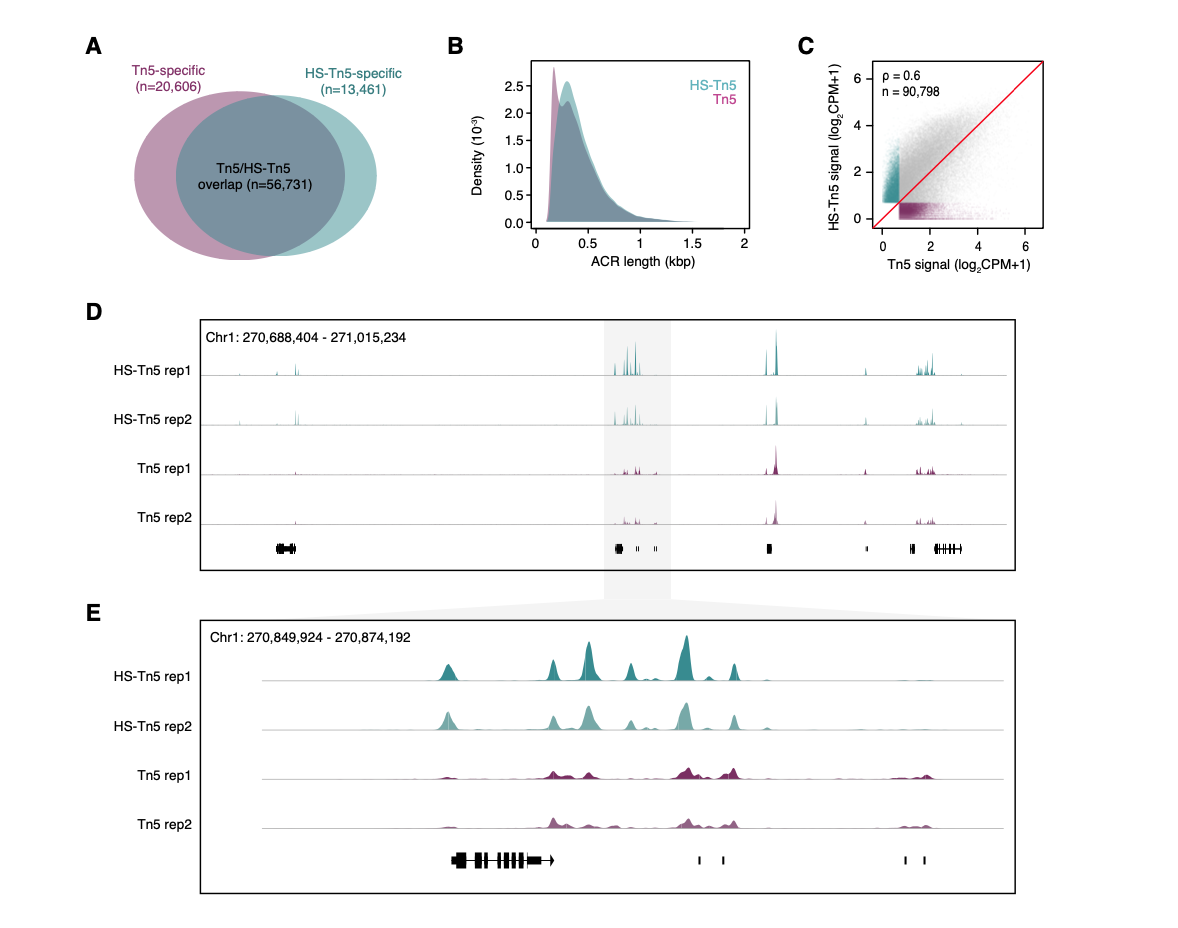

**Supplementary Figure 2. Comparison of TS-Tn5 and Tn5 in *Zea mays* leaves. (A)** Overlapping and specific ACRs identified from TS-Tn5 and Tn5 data sets. **(B)** Distribution of ACR lengths for HS-Tn5 and Tn5. **(C)** Comparison of log2 normalized counts per million (CPM) across the union of ACRs from HS-Tn5 and Tn5. **(D)** and **(E)** Comparison of coverage pile-ups for TS-Tn5 and Tn5.

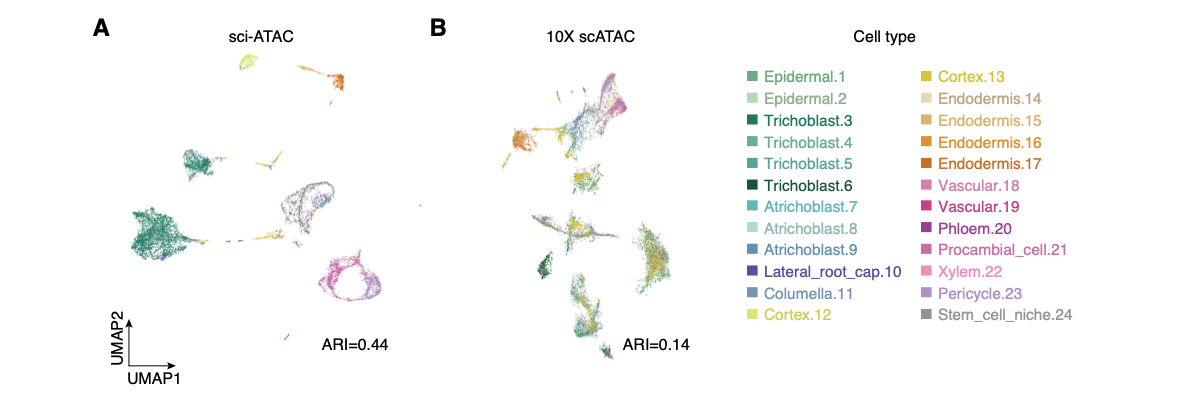

**Supplementary Figure 3.** Comparison of cluster separation, replicate overlap, and concordance with snRNA-seq based cell-type labels. **(A)** sci-ATAC-seq and **(B)** 10x scATAC-seq UMAP colored by snRNA-seq based cell-type labels.

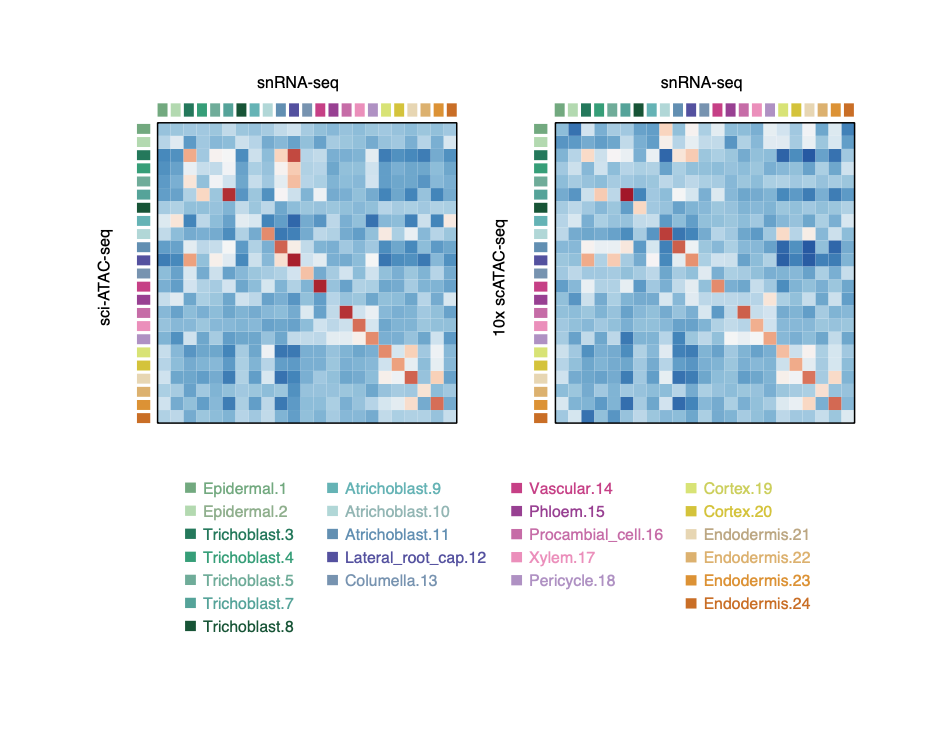

**Supplementary Figure 4.** Pearson correlation coefficients between gene body accessibility and RNA abundance among clusters for sci-ATAC-seq and 10X scATAC-seq.

**Table S1. List of marker genes used for the Arabidopsis root cell type identification.**

| **chr** | **start** | **end** | **geneID** | **name** | **type** | **ref** |
| --- | --- | --- | --- | --- | --- | --- |
| Chr1 | 8030587 | 8033156 | AT1G22710 | SUC2 | Phloem | [10.1007/BF00203657](http://dx.doi.org/10.1007/BF00203657) |
| Chr1 | 18101643 | 18104661 | AT1G48930 | GH9c1 | Hair_cell | 10.1016/j.molp.2021.01.001 |
| Chr1 | 24257216 | 24258473 | AT1G65310 | XTH17 | Non_hair_cell | [10.1111/j.1365-313X.2009.03976.x](https://doi.org/10.1111/j.1365-313x.2009.03976.x) |
| Chr1 | 26027976 | 26030234 | AT1G69240 | AT1G69240 | Hair_cell | [10.1104/pp.109.140905](https://doi.org/10.1104/pp.109.140905) |
| Chr1 | 26288278 | 26293053 | AT1G69830 | AT1G69830 | Columella | [10.1126/science.1146265](https://doi.org/10.1126/science.1146265) |
| Chr1 | 29877228 | 29879279 | AT1G79430 | APL | Phloem | [10.1126/science.1253736](https://doi.org/10.1126/science.1253736) |
| Chr1 | 30036956 | 30041440 | AT1G79840 | GL2 | Non_hair_cell | [10.1038/nature04269](https://doi.org/10.1038/nature04269) |
| Chr2 | 9048131 | 9049404 | AT2G21100 | AT2G21100 | Endodermis | [10.1126/science.1146265](https://doi.org/10.1126/science.1146265) |
| Chr2 | 11708450 | 11710004 | AT2G27370 | CASP3 | Endodermis | [10.1038/nature10070](https://doi.org/10.1038/nature10070) |
| Chr2 | 15644840 | 15647065 | AT2G37260 | TTG2 | Non_hair_cell | [10.1105/tpc.107.052274](https://doi.org/10.1105/tpc.107.052274) |
| Chr3 | 2864782 | 2867230 | AT3G09330 | AT3G09330 | Hair_cell | [10.1126/science.1146265](https://doi.org/10.1126/science.1146265) |
| Chr3 | 3527280 | 3528440 | AT3G11260 | WOX5 | QC | [10.1038/nature05703](https://doi.org/10.1038/nature05703) |
| Chr3 | 3638195 | 3639197 | AT3G11550 | CASP2 | Endodermis | [10.1038/nature10070](https://doi.org/10.1038/nature10070) |
| Chr3 | 19349652 | 19353596 | AT3G52180 | SEX4 | Columella | [10.1126/science.1146265](https://doi.org/10.1126/science.1146265) |
| Chr3 | 20069987 | 20072878 | AT3G54220 | SCR | Endodermis | [10.1038/nature08977](https://doi.org/10.1038/nature08977) |
| Chr3 | 21548484 | 21549572 | AT3G58190 | LBD29 | Pericycle | [10.1093/aob/mcs019](https://doi.org/10.1093/aob/mcs019) |
| Chr4 | 314353 | 317675 | AT4G00750 | AT4G00750 | Columella | [10.1126/science.1146265](https://doi.org/10.1126/science.1146265) |
| Chr4 | 2424164 | 2427769 | AT4G04760 | AT4G04760 | Hair_cell | [10.1371/journal.pone.0055731](https://doi.org/10.1371/journal.pone.0055731) |
| Chr4 | 9655837 | 9656577 | AT4G17215 | AT4G17215 | Endodermis | [10.1371/journal.pgen.1002446](https://doi.org/10.1371/journal.pgen.1002446) |
| Chr4 | 17691687 | 17693859 | AT4G37650 | SHR | Procambial_cell | [10.1038/nature08977](https://doi.org/10.1038/nature08977) |
| Chr5 | 4763372 | 4764823 | AT5G14750 | WER | Non_hair_cell | [10.1016/s0092-8674(00)81536-6](https://doi.org/10.1016/s0092-8674(00)81536-6) |
| Chr5 | 5772568 | 5775417 | AT5G17520 | RCP1 | Root_cap | [10.1073/pnas.96.22.12941](https://doi.org/10.1073/pnas.96.22.12941) |
| Chr5 | 6040919 | 6043014 | AT5G18270 | ANAC087 | Columella | [10.1105/tpc.18.00293](https://doi.org/10.1105/tpc.18.00293) |
| Chr5 | 19248249 | 19249622 | AT5G47450 | TIP2-3 | Pericycle | [10.1186/1471-2229-9-133](https://dx.doi.org/10.1186%2F1471-2229-9-133) |
| Chr5 | 19971884 | 19974530 | AT5G49270 | COBL9 | Hair_cell | [10.1105/tpc.12.10.1961](https://doi.org/10.1105/tpc.12.10.1961) |
| Chr5 | 21649532 | 21651593 | AT5G53370 | AT5G53370 | Cortex | [10.1126/science.1146265](https://doi.org/10.1126/science.1146265) |
| Chr5 | 22407473 | 22410944 | AT5G55250 | IAMT1 | Cortex | [10.1104/pp.18.01482](https://doi.org/10.1104/pp.18.01482) |
| Chr4 | 15863452 | 15868625 | AT4G32880 | HB8 | Xylem | [10.1104/pp.126.2.643](https://doi.org/10.1104/pp.126.2.643) |
| Chr5 | 25050684 | 25051858 | AT5G62380 | VND6 | Xylem | 10.1101/gad.1331305 |
| Chr1 | 27076140 | 27077957 | AT1G71930 | VND7 | Xylem | 10.1101/gad.1331305 |
| Chr2 | 9924933 | 9926372 | AT2G23320 | WRKY15 | Procambial_cell | [10.1105/tpc.19.00689](https://doi.org/10.1105/tpc.19.00689) |
| Chr4 | 14886027 | 14886675 | AT4G30450 | XPP | Pericycle | 10.1038/nature25976 |
| Chr5 | 24724541 | 24727842 | AT5G61480 | PXY | Procambial_cell | [10.1016/j.cub.2007.05.049](https://doi.org/10.1016/j.cub.2007.05.049) |
| Chr5 | 23144980 | 23149568 | AT5G57130 | SMXL5 | Procambial_cell | [10.1016/j.cub.2017.03.014](https://dx.doi.org/10.1016%2Fj.cub.2017.03.014) |
| Chr1 | 11865343 | 11866950 | AT1G32770 | NAC012 | Xylem | [10.1111/j.1365-313X.2007.03109.x](https://doi.org/10.1111/j.1365-313x.2007.03109.x) |
| Chr1 | 17236678  17238053 | AT1G46480 | WOX4 | Procambial_cell | NA | [10.1104/pp.109.149641](https://doi.org/10.1104/pp.109.149641) |
| Chr3 | 9369519 | 9371394 | AT3G25710 | TMO5 | Xylem | [10.1016/j.devcel.2012.12.013](https://doi.org/10.1016/j.devcel.2012.12.013) |
| Chr5 | 24964700 | 24968520 | AT5G62165 | AGL42 | QC | [10.1105/tpc.105.031724](https://dx.doi.org/10.1105%2Ftpc.105.031724) |
| Chr3 | 7300675 | 7303693 | AT3G20840 | PLT1 | QC | [10.1016/j.cell.2004.09.018](https://doi.org/10.1016/j.cell.2004.09.018) |
| Chr1 | 18977284 | 18980618 | AT1G51190 | PLT2 | QC | [10.1016/j.cell.2004.09.018](https://doi.org/10.1016/j.cell.2004.09.018) |
| Chr5 | 3490832 | 3495132 | AT5G11030 | ALF4 | Lateral_root_primordia | [10.1101/gad.9.17.2131](https://doi.org/10.1101/gad.9.17.2131) |
| Chr3 | 1053366 | 1055163 | AT3G04060 | ANAC046 | Columella | [10.1105/tpc.18.00293](https://doi.org/10.1105/tpc.18.00293) |
| Chr5 | 3315529 | 3320202 | AT5G10510 | PLT3 | Lateral_root_primordia | [10.1016/j.cell.2004.09.018](https://doi.org/10.1016/j.cell.2004.09.018) |
| Chr5 | 23253459 | 23256450 | AT5G57390 | PLT5 | Lateral_root_primordia | [10.1016/j.cell.2004.09.018](https://doi.org/10.1016/j.cell.2004.09.018) |
| Chr5 | 4062724 | 4064992 | AT5G12870 | MYB46 | Xylem | 10.1105/tpc.107.053678 |
| Chr4 | 17110747 | 17114141 | AT4G36160 | VND2 | Xylem | [10.1371/journal.pone.0105726](https://dx.doi.org/10.1371%2Fjournal.pone.0105726) |
| Chr1 | 20481690 | 20484541 | AT1G54940 | GUX4 | Hair_cell | [10.1126/science.1146265](https://doi.org/10.1126/science.1146265) |
| Chr1 | 22065617 | 22066973 | AT1G59940 | ARR3 | Passage_cell | 10.1038/nature25976 |
| Chr5 | 25252386 | 25254212 | AT5G62920 | ARR6 | Passage_cell | 10.1038/nature25976 |
| Chr2 | 332967 | 334696 | AT2G01760 | ARR14 | Passage_cell | 10.1038/nature25976 |
| Chr5 | 23334817 | 23336566 | AT5G57620 | MYB36 | Endodermis | 10.1073/pnas.1515576112 |
| Chr5 | 19595657 | 19598492 | AT5G48360 | FH9 | QC | 10.1101/2020.06.29.178863 |
| Chr4 | 13074133 | 13076429 | AT4G25630 | MED36a | Meristem | 10.1101/2020.06.29.178863 |
| Chr3 | 873749 | 876800 | AT3G03620 | DTX24 | Columella | 10.1101/2020.06.29.178863 |
| Chr4 | 10038126 | 10042248 | AT4G18120 | ML3 | Columella | 10.1101/2020.06.29.178863 |
| Chr4 | 11133186 | 11133970 | AT4G20780 | CML42 | Columella | 10.1101/2020.06.29.178863 |
| Chr4 | 17494705 | 17497124 | AT4G37160 | sks15 | Lateral_Root_Cap | 10.1101/2020.06.29.178863 |
| Chr2 | 15034037 | 15036518 | AT2G35770 | scpl28 | Lateral_Root_Cap | 10.1101/2020.06.29.178863 |
| Chr3 | 9496102 | 9497271 | AT3G25950 | AT3G25950 | Lateral_Root_Cap | 10.1101/2020.06.29.178863 |
| Chr5 | 24526847 | 24528084 | AT5G60950 | COBL5 | Non_hair_cell | 10.1101/2020.06.29.178863 |
| Chr3 | 326397 | 327412 | AT3G01970 | WRKY45 | Non_hair_cell | 10.1101/2020.06.29.178863 |
| Chr1 | 9427858 | 9428716 | AT1G27140 | GSTU14 | Hair_cell | 10.1101/2020.06.29.178863 |
| Chr2 | 14740468 | 14743417 | AT2G34940 | VSR5 | Hair_cell | 10.1101/2020.06.29.178863 |
| Chr4 | 13130297 | 13131788 | AT4G25820 | XTH14 | Hair_cell | 10.1101/2020.06.29.178863 |
| Chr5 | 380155 | 380595 | AT5G02000 | AT5G02000 | Cortex | 10.1101/2020.06.29.178863 |
| Chr5 | 25831834 | 25832502 | AT5G64620 | CvIF2 | Cortex | 10.1101/2020.06.29.178863 |
| Chr4 | 6148758 | 6151399 | AT4G09760 | AT4G09760 | Cortex | 10.1101/2020.06.29.178863 |
| Chr2 | 6393332 | 6394938 | AT2G14880 | AT2G14880 | Pericycle | 10.1101/2020.06.29.178863 |
| Chr2 | 18262210 | 18265346 | AT2G44160 | MTHFR2 | Pericycle | 10.1101/2020.06.29.178863 |
| Chr1 | 5578431 | 5580693 | AT1G16310 | MTP10 | Pericycle | 10.1101/2020.06.29.178863 |
| Chr5 | 23851905 | 23855275 | AT5G59090 | SBT4.12 | Phloem | 10.1101/2020.06.29.178863 |
| Chr3 | 1140193 | 1141030 | AT3G04300 | AT3G04300 | Phloem | 10.1101/2020.06.29.178863 |
| Chr2 | 14384896 | 14386669 | AT2G34060 | PER19 | Xylem | 10.1101/2020.06.29.178863 |
| Chr1 | 25861123 | 25863120 | AT1G68810 | bHLH30 | Xylem | 10.1101/2020.06.29.178863 |
| Chr5 | 24167996 | 24170462 | AT5G60020 | LAC17 | Xylem | 10.1101/2020.06.29.178863 |
| Chr5 | 6479400 | 6480446 | AT5G19260 | FAF3 | Procambial_cell | 10.1101/2020.06.29.178863 |
| Chr4 | 8400574 | 8402911 | AT4G14650 | AT4G14650 | Procambial_cell | 10.1101/2020.06.29.178863 |
| Chr1 | 22753715 | 22756300 | AT1G61660 | bHLH112 | Procambial_cell | 10.1101/2020.06.29.178863 |
| Chr2 | 14639323 | 14644355 | AT2G34710 | PHB | Procambial_cell | 10.1016/j.cub.2003.09.035 |
| Chr1 | 24794714 | 24796988 | AT1G66470 | RHD6 | Hair_cell | 10.1016/j.molp.2021.01.001 |
| Chr3 | 20200688 | 20204282 | AT3G54580 | AT3G54580 | Hair_cell | 10.1016/j.molp.2021.01.001 |
| Chr1 | 4276019 | 4277899 | AT1G12560 | EXPA7 | Hair_cell | 10.1016/j.molp.2021.01.001 |
| Chr3 | 23182222 | 23183994 | AT3G62680 | PRP3 | Hair_cell | 10.1016/j.molp.2021.01.001 |
| Chr4 | 279596 | 281199 | AT4G00680 | ADF8 | Hair_cell | 10.1016/j.molp.2021.01.001 |
| Chr4 | 18580838 | 18582085 | AT4G40090 | AGP3 | Hair_cell | 10.1016/j.molp.2021.01.001 |
| Chr5 | 26031184 | 26034389 | AT5G65160 | TPR14 | Hair_cell | 10.1016/j.molp.2021.01.001 |
| Chr4 | 16514827 | 16519534 | AT4G34580 | COW1 | Hair_cell | 10.1016/j.molp.2021.01.001 |
| Chr1 | 2971593 | 2972699 | AT1G09200 | H3.1 | Non_hair_cell | 10.1016/j.molp.2021.01.001 |
| Chr2 | 16792317 | 16793031 | AT2G40205 | AT2G40205 | Non_hair_cell | 10.1016/j.molp.2021.01.001 |
| Chr2 | 18052502 | 18054387 | AT2G43480 | AT2G43480 | Non_hair_cell | 10.1016/j.molp.2021.01.001 |
| Chr1 | 9889125 | 9891493 | AT1G28290 | AGP31 | Non_hair_cell | 10.1016/j.molp.2021.01.001 |
| Chr1 | 28756021 | 28759297 | AT1G76620 | AT1G76620 | Non_hair_cell | 10.1016/j.molp.2021.01.001 |
| Chr4 | 299359 | 305008 | AT4G00730 | ANL2 | Non_hair_cell | 10.1016/j.molp.2021.01.001 |
| Chr1 | 12233269 | 12236476 | AT1G33750 | AT1G33750 | Non_hair_cell | 10.1016/j.molp.2021.01.001 |
| Chr4 | 12934062 | 12935373 | AT4G25250 | PMEI4 | Non_hair_cell | 10.1016/j.molp.2021.01.001 |
| Chr1 | 11475307 | 11478194 | AT1G31950 | AT1G31950 | Non_hair_cell | 10.1016/j.molp.2021.01.001 |
| Chr4 | 8721487 | 8727169 | AT4G15290 | ATCSLB05 | Non_hair_cell | 10.1016/j.molp.2021.01.001 |
| Chr5 | 22074783 | 22076777 | AT5G54370 | AT5G54370 | Columella | 10.1016/j.molp.2021.01.001 |
| Chr1 | 3157001 | 3159148 | AT1G09750 | AED3 | Cortex | 10.1016/j.molp.2021.01.001 |
| Chr3 | 9639199 | 9641366 | AT3G26300 | CYP71B34 | Cortex | 10.1016/j.molp.2021.01.001 |
| Chr1 | 23136369 | 23137691 | AT1G62510 | AT1G62510 | Cortex | 10.1016/j.molp.2021.01.001 |
| Chr5 | 6282381 | 6286707 | AT5G18840 | AT5G18840 | Cortex | 10.1016/j.molp.2021.01.001 |
| Chr3 | 7626757 | 7629651 | AT3G21670 | NPF6.4 | Cortex | 10.1016/j.molp.2021.01.001 |
| Chr1 | 5602373 | 5604663 | AT1G16390 | OCT3 | Endodermis | 10.1016/j.molp.2021.01.001 |
| Chr1 | 22723501 | 22726574 | AT1G61590 | PBL15 | Endodermis | 10.1016/j.molp.2021.01.001 |
| Chr2 | 16776890 | 16779408 | AT2G40160 | TBL30 | Endodermis | 10.1016/j.molp.2021.01.001 |
| Chr2 | 19685066 | 19686493 | AT2G48130 | AT2G48130 | Endodermis | 10.1016/j.molp.2021.01.001 |
| Chr3 | 8008034 | 8009590 | AT3G22620 | AT3G22620 | Endodermis | 10.1016/j.molp.2021.01.001 |
| Chr4 | 6869453 | 6871661 | AT4G11290 | AT4G11290 | Endodermis | 10.1016/j.molp.2021.01.001 |
| Chr1 | 1529267 | 1531439 | AT1G05260 | PER3 | Endodermis | 10.1016/j.molp.2021.01.001 |
| Chr3 | 20471291 | 20472889 | AT3G55230 | AT3G55230 | Endodermis | 10.1016/j.molp.2021.01.001 |
| Chr1 | 17001617 | 17003688 | AT1G44970 | PER9 | Endodermis | 10.1016/j.molp.2021.01.001 |
| Chr2 | 6649130 | 6652010 | AT2G15300 | AT2G15300 | Endodermis | 10.1016/j.molp.2021.01.001 |
| Chr2 | 539499 | 540634 | AT2G02130 | PDF2.3 | Columella | 10.1016/j.molp.2021.01.001 |
| Chr3 | 21941865 | 21943071 | AT3G59370 | AT3G59370 | Columella | 10.1016/j.molp.2021.01.001 |
| Chr1 | 26742554 | 26746395 | AT1G70940 | PIN3 | Columella | 10.1016/j.molp.2021.01.001 |
| Chr1 | 7251674 | 7253731 | AT1G20850 | XCP2 | Xylem | 10.1016/j.molp.2021.01.001 |
| Chr3 | 2576794 | 2578746 | AT3G08500 | MYB83 | Xylem | 10.1016/j.molp.2021.01.001 |
| Chr4 | 16809982 | 16812009 | AT4G35350 | XCP1 | Xylem | 10.1016/j.molp.2021.01.001 |
| Chr5 | 5084204 | 5086792 | AT5G15630 | COBL4 | Xylem | 10.1016/j.molp.2021.01.001 |
| Chr3 | 3107275 | 3108677 | AT3G10080 | AT3G10080 | Xylem | 10.1016/j.molp.2021.01.001 |
| Chr5 | 594179 | 595367 | AT5G02640 | AT5G02640 | Xylem | 10.1016/j.molp.2021.01.001 |
| Chr2 | 8144412 | 8146265 | AT2G18800 | XTH21 | Pericycle | 10.1016/j.molp.2021.01.001 |
| Chr5 | 21293687 | 21296704 | AT5G52470 | FIB1 | Pericycle | 10.1016/j.molp.2021.01.001 |
| Chr2 | 6653291 | 6655501 | AT2G15310 | ARFB1A | Phloem | 10.1016/j.molp.2021.01.001 |
| Chr3 | 832429 | 834350 | AT3G03500 | AT3G03500 | Phloem | 10.1016/j.molp.2021.01.001 |
| Chr1 | 13363003 | 1336556 | AT1G35910 | TPPD | Pericycle | 10.1016/j.molp.2021.01.001 |
| Chr4 | 12223453 | 12225323 | AT4G23410 | TET5 | Pericycle | 10.1016/j.molp.2021.01.001 |
| Chr1 | 29940850 | 29944102 | AT1G79580 | SMB | Root_cap | 10.1016/j.molp.2021.01.001 |
| Chr4 | 2223922 | 2227874 | AT4G04460 | PASPA3 | Root_cap | 10.1016/j.molp.2021.01.001 |
| Chr2 | 13835734 | 13839513 | AT2G32610 | CSLB01 | Lateral_Root_Cap | 10.1016/j.molp.2021.01.001 |
| Chr1 | 18171095 | 18172450 | AT1G49110 | AT1G49110 | QC | 10.1016/j.molp.2021.01.001 |
| Chr2 | 1170636 | 1172363 | AT2G03830 | RGF8 | QC | 10.1016/j.molp.2021.01.001 |
| Chr3 | 9545898 | 9549453 | AT3G26120 | TEL1 | QC | 10.1016/j.molp.2021.01.001 |
| Chr5 | 5742542 | 5746068 | AT5G17430 | BBM | QC | 10.1016/j.molp.2021.01.001 |
| Chr5 | 8017192 | 8019526 | AT5G23780 | DUF9 | QC | 10.1016/j.molp.2021.01.001 |
| Chr5 | 24466531 | 24467679 | AT5G60810 | RGF1 | QC | 10.1016/j.molp.2021.01.001 |
| Chr1 | 25769576 | 25773043 | AT1G68640 | PAN | QC | 10.1016/j.molp.2021.01.001 |
| Chr1 | 28017584 | 28020549 | AT1G74560 | NRP1 | Lateral_root_primordia | 10.1101/2020.10.02.324327 |
| Chr5 | 3987375 | 3990333 | AT5G12330 | LRP1 | Lateral_root_primordia | 10.1101/2020.10.02.324327 |

**Table S2 | List of index Tn5 adapter sequences.**

| adapter | **sequence** | **index** |
| --- | --- | --- |
| Tn5-ME-A1 | GGCGGTAGGCGTGatacgAGATGTGTATAAGAGACAG | **ATACG** |
| Tn5-ME-A2 | GGCGGTAGGCGTGagcgaAGATGTGTATAAGAGACAG | **AGCGA** |
| Tn5-ME-A3 | GGCGGTAGGCGTGctaagAGATGTGTATAAGAGACAG | **CTAAG** |
| Tn5-ME-A4 | GGCGGTAGGCGTGtgtcgAGATGTGTATAAGAGACAG | **TGTCG** |
| Tn5-ME-A5 | GGCGGTAGGCGTGgtagtAGATGTGTATAAGAGACAG | **GTAGT** |
| Tn5-ME-A6 | GGCGGTAGGCGTGgtataAGATGTGTATAAGAGACAG | **GTATA** |
| Tn5-ME-A7 | GGCGGTAGGCGTGattccAGATGTGTATAAGAGACAG | **ATTCC** |
| Tn5-ME-A8 | GGCGGTAGGCGTGgacttAGATGTGTATAAGAGACAG | **GACTT** |
| Tn5-ME-A9 | GGCGGTAGGCGTGttactAGATGTGTATAAGAGACAG | **TTACT** |
| Tn5-ME-A10 | GGCGGTAGGCGTGcaattAGATGTGTATAAGAGACAG | **CAATT** |
| Tn5-ME-A11 | GGCGGTAGGCGTGcatctAGATGTGTATAAGAGACAG | **CATCT** |
| Tn5-ME-A12 | GGCGGTAGGCGTGtcgcgAGATGTGTATAAGAGACAG | **TCGCG** |
| Tn5-ME-A13 | GGCGGTAGGCGTGtgtttAGATGTGTATAAGAGACAG | **TGTTT** |
| Tn5-ME-A14 | GGCGGTAGGCGTGtccctAGATGTGTATAAGAGACAG | **TCCCT** |
| Tn5-ME-A15 | GGCGGTAGGCGTGggcagAGATGTGTATAAGAGACAG | **GGCAG** |
| Tn5-ME-A16 | GGCGGTAGGCGTGcaaaaAGATGTGTATAAGAGACAG | **CAAAA** |
| Tn5-ME-A17 | GGCGGTAGGCGTGgtgagAGATGTGTATAAGAGACAG | **GTGAG** |
| Tn5-ME-A18 | GGCGGTAGGCGTGtggatAGATGTGTATAAGAGACAG | **TGGAT** |
| Tn5-ME-A19 | GGCGGTAGGCGTGccgagAGATGTGTATAAGAGACAG | **CCGAG** |
| Tn5-ME-A20 | GGCGGTAGGCGTGactcgAGATGTGTATAAGAGACAG | **ACTCG** |
| Tn5-ME-A21 | GGCGGTAGGCGTGgtcctAGATGTGTATAAGAGACAG | **GTCCT** |
| Tn5-ME-A22 | GGCGGTAGGCGTGtaagaAGATGTGTATAAGAGACAG | **TAAGA** |
| Tn5-ME-A23 | GGCGGTAGGCGTGaccccAGATGTGTATAAGAGACAG | **ACCCC** |
| Tn5-ME-A24 | GGCGGTAGGCGTGaaacaAGATGTGTATAAGAGACAG | **AAACA** |
| Tn5-ME-B1 | CCAACACCCGTGCGtcggaAGATGTGTATAAGAGACAG | **TCGGA** |
| Tn5-ME-B2 | CCAACACCCGTGCGgtttcAGATGTGTATAAGAGACAG | **GTTTC** |
| Tn5-ME-B3 | CCAACACCCGTGCGagcttAGATGTGTATAAGAGACAG | **AGCTT** |
| Tn5-ME-B4 | CCAACACCCGTGCGtcattAGATGTGTATAAGAGACAG | **TCATT** |
| Tn5-ME-B5 | CCAACACCCGTGCGgctccAGATGTGTATAAGAGACAG | **GCTCC** |
| Tn5-ME-B6 | CCAACACCCGTGCGactaaAGATGTGTATAAGAGACAG | **ACTAA** |
| Tn5-ME-B7 | CCAACACCCGTGCGcgaggAGATGTGTATAAGAGACAG | **CGAGG** |
| Tn5-ME-B8 | CCAACACCCGTGCGaccggAGATGTGTATAAGAGACAG | **ACCGG** |
| Tn5-ME-B9 | CCAACACCCGTGCGcggttAGATGTGTATAAGAGACAG | **CGGTT** |
| Tn5-ME-B10 | CCAACACCCGTGCGggaacAGATGTGTATAAGAGACAG | **GGAAC** |
| Tn5-ME-B11 | CCAACACCCGTGCGgatgaAGATGTGTATAAGAGACAG | **GATGA** |
| Tn5-ME-B12 | CCAACACCCGTGCGaacatAGATGTGTATAAGAGACAG | **AACAT** |
| Tn5-ME-B13 | CCAACACCCGTGCGcgcacAGATGTGTATAAGAGACAG | **CGCAC** |
| Tn5-ME-B14 | CCAACACCCGTGCGtgaccAGATGTGTATAAGAGACAG | **TGACC** |
| Tn5-ME-B15 | CCAACACCCGTGCGcgcgtAGATGTGTATAAGAGACAG | **CGCGT** |
| Tn5-ME-B16 | CCAACACCCGTGCGttcccAGATGTGTATAAGAGACAG | **TTCCC** |

**Table S3. List of PCR primers.**

| **Row index**  **primer** | **Sequence** | **Index** |
| --- | --- | --- |
| A | ACACTCTTTCCCTACACGACGCTCTTCCGATCTGATCAGGGCGGTAGGCGTG | **GATCAG** |
| B | ACACTCTTTCCCTACACGACGCTCTTCCGATCTTAGCTTGGCGGTAGGCGTG | **TAGCTT** |
| C | ACACTCTTTCCCTACACGACGCTCTTCCGATCTGGCTACGGCGGTAGGCGTG | **GGCTAC** |
| D | ACACTCTTTCCCTACACGACGCTCTTCCGATCTCTTGTAGGCGGTAGGCGTG | **CTTGTA** |
| E | ACACTCTTTCCCTACACGACGCTCTTCCGATCTAGTCAAGGCGGTAGGCGTG | **AGTCAA** |
| F | ACACTCTTTCCCTACACGACGCTCTTCCGATCTAGTTCCGGCGGTAGGCGTG | **AGTTCC** |
| G | ACACTCTTTCCCTACACGACGCTCTTCCGATCTATGTCAGGCGGTAGGCGTG | **ATGTCA** |
| H | ACACTCTTTCCCTACACGACGCTCTTCCGATCTCCGTCCGGCGGTAGGCGTG | **CCGTCC** |
| **Column index**  **primer** | **Sequence** | **index** |
| 1 | GTGACTGGAGTTCAGACGTGTGCTCTTCCGATCTGTAGAGCCAACACCCGTGCG | **GTAGAG** |
| 2 | GTGACTGGAGTTCAGACGTGTGCTCTTCCGATCTGTCCGCCCAACACCCGTGCG | **GTCCGC** |
| 3 | GTGACTGGAGTTCAGACGTGTGCTCTTCCGATCTGTGAAACCAACACCCGTGCG | **GTGAAA** |
| 4 | GTGACTGGAGTTCAGACGTGTGCTCTTCCGATCTGTGGCCCCAACACCCGTGCG | **GTGGCC** |
| 5 | GTGACTGGAGTTCAGACGTGTGCTCTTCCGATCTGTTTCGCCAACACCCGTGCG | **GTTTCG** |
| 6 | GTGACTGGAGTTCAGACGTGTGCTCTTCCGATCTCGTACGCCAACACCCGTGCG | **CGTACG** |
| 7 | GTGACTGGAGTTCAGACGTGTGCTCTTCCGATCTGAGTGGCCAACACCCGTGCG | **GAGTGG** |
| 8 | GTGACTGGAGTTCAGACGTGTGCTCTTCCGATCTGGTAGCCCAACACCCGTGCG | **GGTAGC** |
| 9 | GTGACTGGAGTTCAGACGTGTGCTCTTCCGATCTACTGATCCAACACCCGTGCG | **ACTGAT** |
| 10 | GTGACTGGAGTTCAGACGTGTGCTCTTCCGATCTATGAGCCCAACACCCGTGCG | **ATGAGC** |
| 11 | GTGACTGGAGTTCAGACGTGTGCTCTTCCGATCTATTCCTCCAACACCCGTGCG | **ATTCCT** |
| 12 | GTGACTGGAGTTCAGACGTGTGCTCTTCCGATCTCAAAAGCCAACACCCGTGCG | **CAAAAG** |
| **Illumina index**  **primer** | **Sequence** | **index** |
| N518 | AATGATACGGCGACCACCGAGATCTACACCGTCTAATACACTCTTTCCCTACACGACGCTCTTCCGATC*T | **ATTAGACG** |
| N519 | AATGATACGGCGACCACCGAGATCTACACTGGTACCCACACTCTTTCCCTACACGACGCTCTTCCGATC*T | **GGGTACCA** |
| N520 | AATGATACGGCGACCACCGAGATCTACACAACATGTGACACTCTTTCCCTACACGACGCTCTTCCGATC*T | **CACATGTT** |
| N521 | AATGATACGGCGACCACCGAGATCTACACGCAGATGAACACTCTTTCCCTACACGACGCTCTTCCGATC*T | **TCATCTGC** |
| N701 | CAAGCAGAAGACGGCATACGAGATTCGCCTTAGTGACTGGAGTTCAGACGTGTGCTCTTCCGATC*T | **TAAGGCGA** |
| N702 | CAAGCAGAAGACGGCATACGAGATCTAGTACGGTGACTGGAGTTCAGACGTGTGCTCTTCCGATC*T | **CGTACTAG** |
| N703 | CAAGCAGAAGACGGCATACGAGATTTCTGCCTGTGACTGGAGTTCAGACGTGTGCTCTTCCGATC*T | **AGGCAGAA** |
| N704 | CAAGCAGAAGACGGCATACGAGATGCTCAGGAGTGACTGGAGTTCAGACGTGTGCTCTTCCGATC*T | **TCCTGAGC** |
